# An opponent striatal circuit for distributional reinforcement learning

**DOI:** 10.1101/2024.01.02.573966

**Authors:** Adam S. Lowet, Qiao Zheng, Melissa Meng, Sara Matias, Jan Drugowitsch, Naoshige Uchida

## Abstract

Machine learning research has achieved large performance gains on a wide range of tasks by expanding the learning target from mean rewards to entire probability distributions of rewards — an approach known as distributional reinforcement learning (RL)^1^. The mesolimbic dopamine system is thought to underlie RL in the mammalian brain by updating a representation of mean value in the striatum^2,3^, but little is known about whether, where, and how neurons in this circuit encode information about higher-order moments of reward distributions^4^. To fill this gap, we used high-density probes (Neuropixels) to acutely record striatal activity from well-trained, water-restricted mice performing a classical conditioning task in which reward mean, reward variance, and stimulus identity were independently manipulated. In contrast to traditional RL accounts, we found robust evidence for abstract encoding of variance in the striatum. Remarkably, chronic ablation of dopamine inputs disorganized these distributional representations in the striatum without interfering with mean value coding. Two-photon calcium imaging and optogenetics revealed that the two major classes of striatal medium spiny neurons — D1 and D2 MSNs — contributed to this code by preferentially encoding the right and left tails of the reward distribution, respectively. We synthesize these findings into a new model of the striatum and mesolimbic dopamine that harnesses the opponency between D1 and D2 MSNs^5–15^ to reap the computational benefits of distributional RL.

## Main Text

Midbrain dopamine neurons and their primary target, the striatum, constitute an evolutionarily ancient^16^ neural circuit that is critical for motivated behaviors^17,18^. Computationally, dopamine has long been thought to signal reward prediction error (RPE)^2,19,20^, reminiscent of the teaching signals used in many reinforcement learning (RL) algorithms^21^. Consistent with this idea, dopamine is also known to modulate plasticity of certain corticostriatal synapses in roughly the manner predicted by RL theory^22^, allowing neurons in the striatum to learn a representation of average anticipated reward^23–28^, often called “value”.

Despite the simplicity and popularity of this model, it leaves many aspects of the mesolimbic circuit unexplained. For one, value representations reside not only in the striatum but throughout the entire brain^29–35^, and are enriched in neurons projecting *to* the striatum^36,37^. Second, the striatum is far from uniform, containing a variety of interneuron subtypes as well as D1 and D2 medium spiny neurons (MSNs), whose projection patterns differ^38^ and whose plasticity is modulated in opposite directions by dopamine^22,39,40^. These differences at the receptor level translate to opposite coding properties^5,6^ and effects on behavior^7–15^, but interpreting their distinct roles is complicated by the fact that these two populations often co-activate^41–45^. Third, dopamine activity is much more complex than a simple scalar RPE, varying both qualitatively across dopamine projection systems^46–49^ and quantitatively within systems^4,50,51^. Whether such diversity is cause to revise RPE-based accounts of dopamine^4,52,53^ or discard them altogether^54,55^ is currently the subject of intense debate.

In parallel to these questions about the neuronal representation of value, the striatum — particularly the ventral striatum (VS, also referred to as the nucleus accumbens) — has long been associated with decision-making under risk. VS lesions^56–58^ and dopaminergic drugs^59,60^ can both impair risky decision-making, with some groups suggesting a particular role for VS D2 MSNs^61,62^. Aberrant processing of risk, in turn, is thought to underlie many diseases associated with these circuits, particularly addiction^63–65^. Given this, it is perhaps surprising that, with a few exceptions^66^, conventional RL models of the basal ganglia ignore the role of risk, and most theoretical investigations of uncertainty focus on sensory noise rather than intrinsic, irreducible environmental stochasticity^67–70^.

Borrowing from tremendous successes and popularity in machine learning^71–73^, it was recently proposed^4^ that the residual heterogeneity within RPE-coding dopamine neurons^74,75^ resembles the predictions of a particular algorithm known as Expectile Distributional RL (EDRL)^76^. This algorithm not only unifies the learning of value and risk (and potentially higher-order moments of reward distributions) within the same biologically-plausible framework but also provides novel computational advantages — even with risk-neutral settings — related to representation learning in deep neural networks^77,78^ and, potentially, directed exploration^79–82^. However, alternative accounts of the same dopamine data have since been put forward^83^, including some that question the very existence of a probabilistic value code^84,85^.

Here, we provide the first direct evidence for distributional RL in the mammalian brain by demonstrating that the striatum, and particularly VS, encodes not just mean value but also reward variance. We combine our observations with well-established features of the anatomy and physiology of the basal ganglia to construct a new computational model of reward distribution learning in the striatum. The proposed model brings together diverse dopamine inputs^4^ and asymmetric plasticity rules^22,39,40^ to enable a biological implementation of EDRL. Our model makes several new experimental predictions about the representational geometry of the striatal population code and its dependence on intact dopamine inputs, which we confirm using Neuropixels recordings and dopamine lesions. Moreover, it suggests a way to unify the opponent yet concurrent and non-redundant contributions of D1 and D2 MSNs to behavior via their coding of the right and left tails of the reward distribution, respectively. We validate this view using cell type-specific two-photon calcium imaging and optogenetic manipulations. Together, this study improves our understanding of the computational principles underlying the brain’s reward circuitry and tightens the bonds between natural and artificial intelligence.

## A behavioral task to investigate distributional RL

Representations of reward variance have been previously observed in a variety of cortical^86–88^ and subcortical^89–91^ regions, but not in the striatum. To determine whether striatal neurons encode reward variance while remaining agnostic to its representational format, we designed a classical conditioning task in which water-restricted mice were trained to associate random odor cues with probability distributions over stochastic reward magnitudes (Fig. 1a). Three different probability distributions (Fig. 1b) were used: Nothing (100% chance of 0 μL reward), Fixed (100% chance of 4 μL reward), and Variable (50/50% chance of 2/6 μL reward). Fixed and Variable distributions shared the same mean but had a different variance. Thus, distributional RL predicts systematic differences in their underlying neural representations, whereas traditional RL — assuming risk neutrality — does not. To ensure any such differences did not reflect idiosyncratic odor preferences, two unique odors predicted each of the three distributions, allowing us to compare odor representations both across-and within-distributions.

**Fig. 1.**
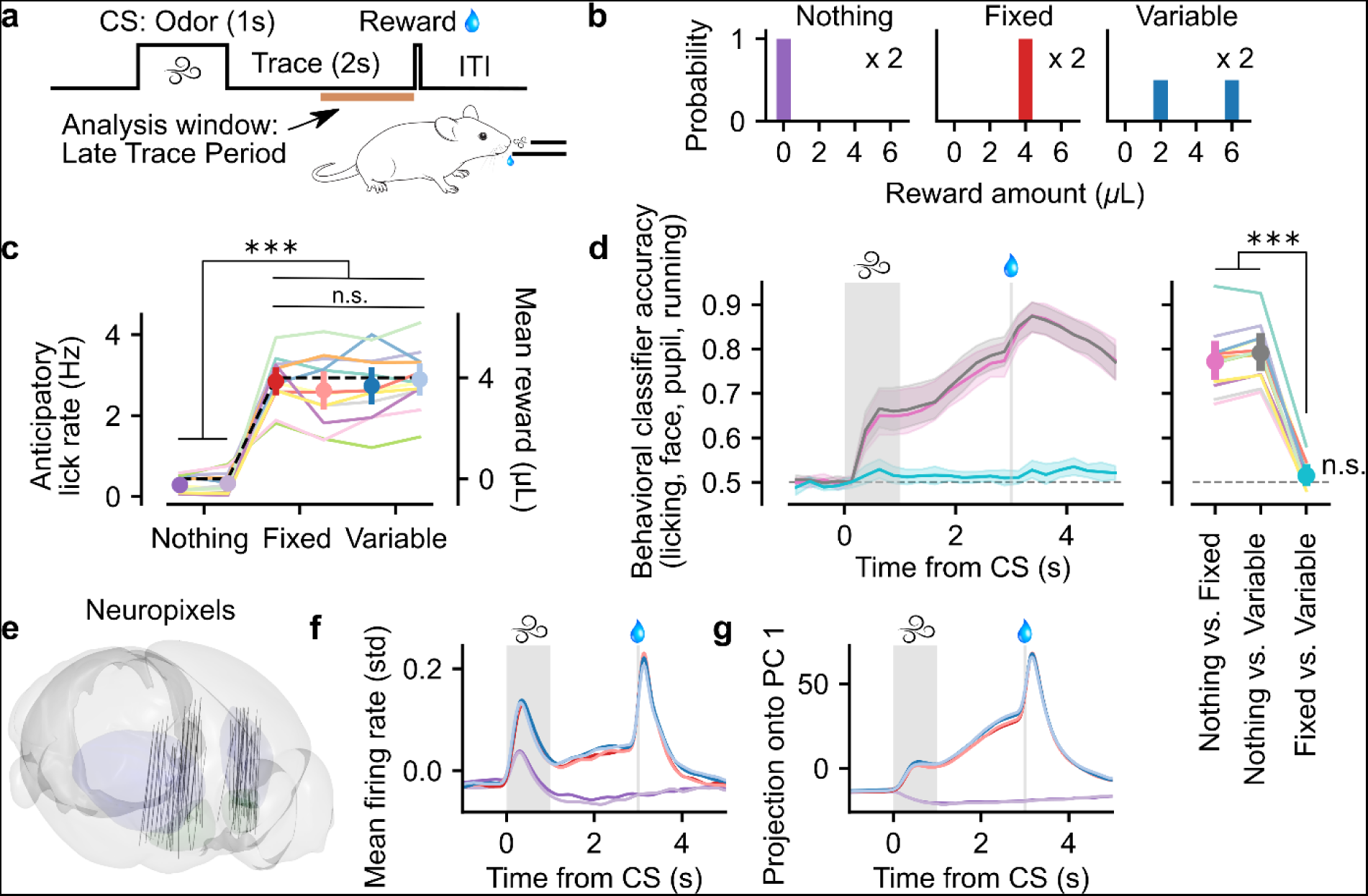
**| A classical conditioning task and recording setup to investigate distributional reinforcement learning. a**, Water-restricted, head-fixed mice were trained to associate odors with stochastic rewards following a brief (2 s) trace period. When not otherwise specified, behavioral and neural activity were analyzed in the final second of the trace period (“Late Trace” period) in order to assess reward anticipation. Odor-reward distribution mappings were randomized across mice. CS, conditioned stimulus. ITI, inter-trial interval. **b,** Probability distributions over reward amounts that were paired with odors. Each distribution was associated with two distinct odors, for a total of six odors, in order to distinguish stimulus information from distributional content. Furthermore, two distributions (Fixed and Variable) had the same mean of 4 μL, but different variance. **c**, Anticipatory lick rates for each trial type, computed during the Late Trace period (Nothing 1 or Nothing 2: *p* < 0.001 versus Fixed 1, Fixed 2, Variable 1, and Variable 2; Fixed 1: *p* = 0.502, 0.925, 0.419 versus Fixed 2, Variable 1, and Variable 2, respectively). **d**, Cross-validated classification accuracy of a linear kernel Support Vector Machine trained on licking, pupil area, whisking, running, and singular value decomposition of behavioral videos (Extended Data Fig. 1). The data associated with the two odors corresponding to the same distribution were pooled and then split into training and validation sets. *Left*, behavioral classifier accuracy across time. Predictors were aggregated within 250 ms, non-overlapping bins. Shaded regions denote 95% confidence intervals across mice. Pink, Nothing vs. Fixed; Grey, Nothing vs. Variable; Cyan, Fixed vs. Variable. *Right*, quantification of behavioral classifier accuracy when trained separately on the entire Late Trace period (Fixed vs. Variable: *p* < 0.001 versus Nothing vs. Fixed and Nothing vs. Variable, *p* = 0.053 compared to chance level of 50%). **e,** Reconstructed Neuropixels probe trajectories, aligned to the Allen Mouse Brain Common Coordinate Framework. **f**, Individual neurons’ firing rates were z-scored across time, aligned to stimulus onset, averaged for each trial type, and then averaged across neurons. Color code as in **c**. Average firing rates correlate with mean reward. **g**, Trial type averages for each neuron were concatenated, and the first principal component was extracted and plotted across neurons. Color code as in **c**. For Figs. 1–3 and 6, asterisks represent the result of Linear Mixed Effects model across sessions with a random intercept for each mouse, and, if applicable, a random slope for each mouse as a function of grouping (e.g. Across-vs. Within-distribution): ∗∗∗, *p* < 0.001; ∗∗, *p* < 0.01; ∗, *p* < 0.05; n.s., not significant at *α* = 0.05. Asterisks over lines connecting different groupings indicate significant differences between groups, while asterisks without corresponding lines indicate that the group is significantly different from chance, indicated by the dashed grey line. The shaded region from 0 to 1 s represents the interval of odor delivery, and the vertical line at 3 s indicates reward timing. For Figures 1–4, pastel colors in the background show averages across sessions within mice, while dots with error bars in the foreground denote means and 95% confidence intervals across mice. Differences were taken within-session.

Crucially, while animals’ anticipatory licking revealed a clear preference for Rewarded (Fixed and Variable) over Unrewarded odors, it did not differ between the Fixed and Variable distributions (Fig. 1c; here and elsewhere, we plot each mouse’s mean across sessions as a colored line for clarity, but the statistical tests disaggregate sessions using a Linear Mixed Effects model with mouse-level random effects; see Methods). Additional behavioral data, including face motion, whisking, pupil area, and running^92^, also did not support reliably distinguishing Fixed from Variable trials (Fig. 1d and Extended Data Fig. 1a-b). The meager, non-significant classification ability that may have existed was orthogonal to the regression weight vector trained to predict value from all trial types (Extended Data Fig. 1d-e). This implies that any ability to decode these trial types from neural data must be due to the associated probability distributions and not to differential valuation or motor behavior.

## Striatum represents both mean and variance

Next, we used high-density electrophysiological probes (Neuropixels) to acutely record activity from across a broad swathe of the anterior striatum (Fig. 1e and Extended Data Fig. 2a; *N* = 12 mice, *n* = 71 sessions, 13,997 neurons). Consistent with prior work^23–28^, we found that both the average firing rate of all neurons (Fig. 1f and Extended Data Fig. 2b) and the time course of trial type-averaged activity projected onto the first principal component (PC; Fig. 1g) cleanly separated Rewarded from Unrewarded odors. Furthermore, a substantial fraction of the activity of individual neurons within our 1 s analysis window just before reward delivery correlated significantly with expected reward, allowing us to reliably predict mean value from neural (pseudo-) population activity across all striatal subregions (Extended Data Fig. 2c-e). Other striatal neurons correlated significantly with reward prediction error during the reward period^93^, but these formed a smaller and mostly independent subset (Extended Data Fig. 2f-h).

**Fig. 2.**
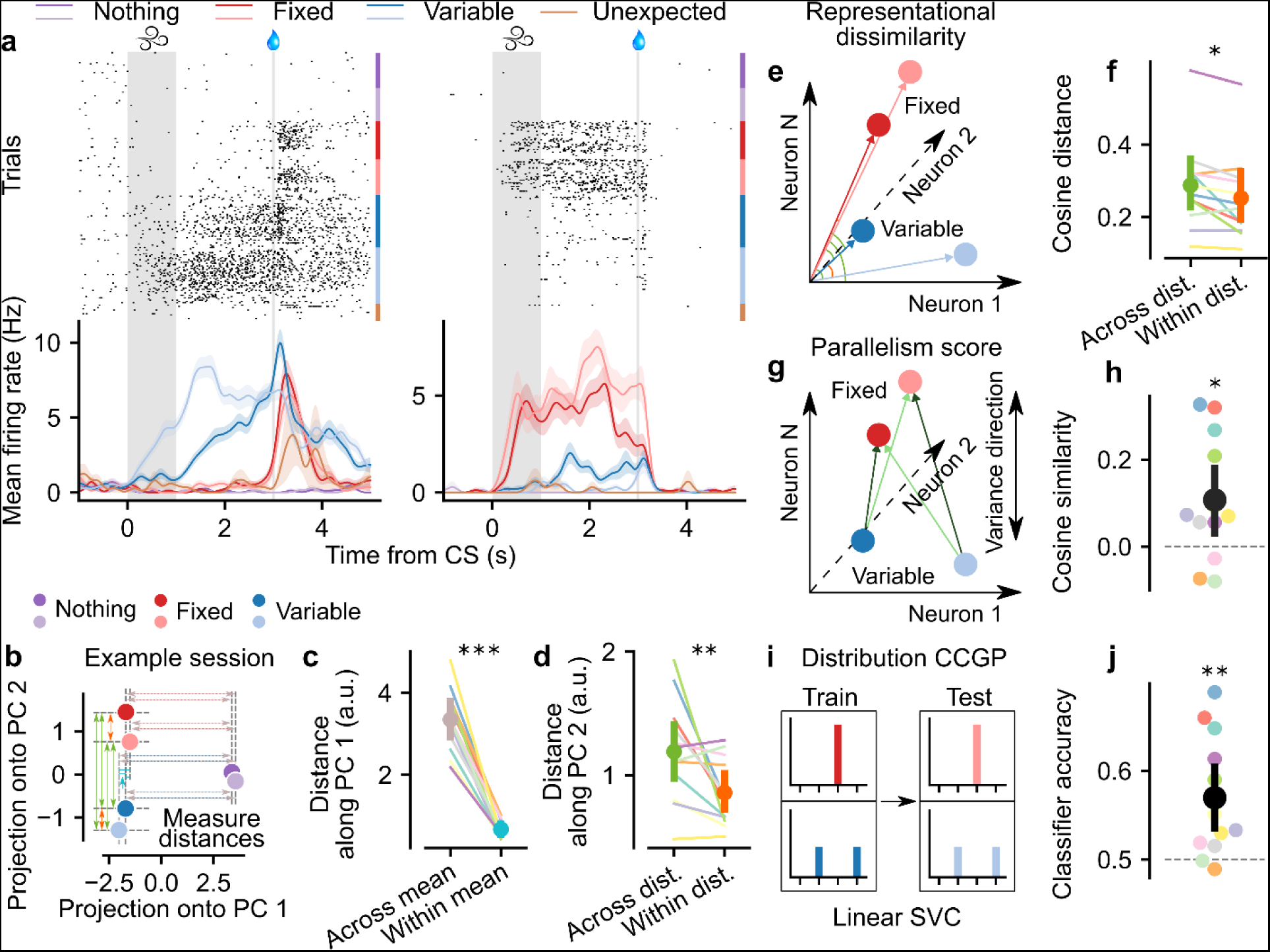
**| Distributional coding across the striatum. a**, Example peri-stimulus time histograms (PSTHs) of two simultaneously-recorded neurons in the ventromedial striatum. *Top*, spike rasters, aligned to odor onset and sorted by trial type. *Bottom*, mean ± s.e.m. firing to each trial type, after smoothing the entire session’s spike train with a Gaussian kernel (s.d. = 100 ms). While both neurons tend to increase on average to rewarded odors, the neuron on the left prefers Variable odors, while the one on the right prefers Fixed odors, and tend to do so consistently for different odors associated with the same distribution. **b**, Firing rate during the Late Trace period, averaged across trials of each type, was projected into two dimensions using principal component analysis (PCA) independently for each session. We then measured the distances between trial types along each PC, as shown by the arrows. Color code as in **a**. **c**, Euclidean distance along PC 1 was significantly greater for across-mean pairs (Nothing vs. Rewarded) than within-mean pairs (Fixed vs. Variable; *p* < 0.001). **d**, Euclidean distance along PC 2 was significantly greater for across-distribution pairs (Fixed vs. Variable) than within-distribution pairs (Fixed 1 vs. Fixed 2 or Variable 1 vs. Variable 2; *p* = 0.006). **e**, Schematic illustrating representational dissimilarity analysis (RDA). The population vector corresponding to each trial type was computed independently for each session. We then computed the cosine distances between across-distribution and within-distribution pairs, shown by the green and orange arcs. **f**, Quantification of cosine distances (Across-vs. Within-distribution: *p* = 0.029). **g**, Schematic illustrating parallelism score. We computed the difference vector between each Fixed and Variable trial type for each session independently. Parallelism score is defined as the cosine similarity between each non-overlapping pair of vectors, averaged over the two possible combinations (dark green and light green). **h**, Quantification of parallelism score (*p* = 0.015 compared to chance level of 0). **i**, Schematic illustrating computation of cross-condition generalization performance (CCGP). Linear support vector classifiers (SVCs) were trained to discriminate one Fixed and one Variable odor and then tested on the held-out Fixed vs. Variable pair. This was then repeated and averaged over all four possible combinations of training and test sets. **j**, Quantification of CCGP (*p* = 0.001 compared to chance level of 50%).

However, not all neurons obeyed this simple pattern seen at the level of population averages. Some single neurons consistently preferred Variable odors, while others — even when recorded simultaneously — preferred Fixed (Fig. 2a). Such neurons fired similarly to *both instances* of the Fixed and Variable odors, suggesting that they abstracted over odor-specific details to instead encode information about variance — even as the population as a whole contained ample odor information (Extended Data Fig. 3a-e).

**Fig. 3.**
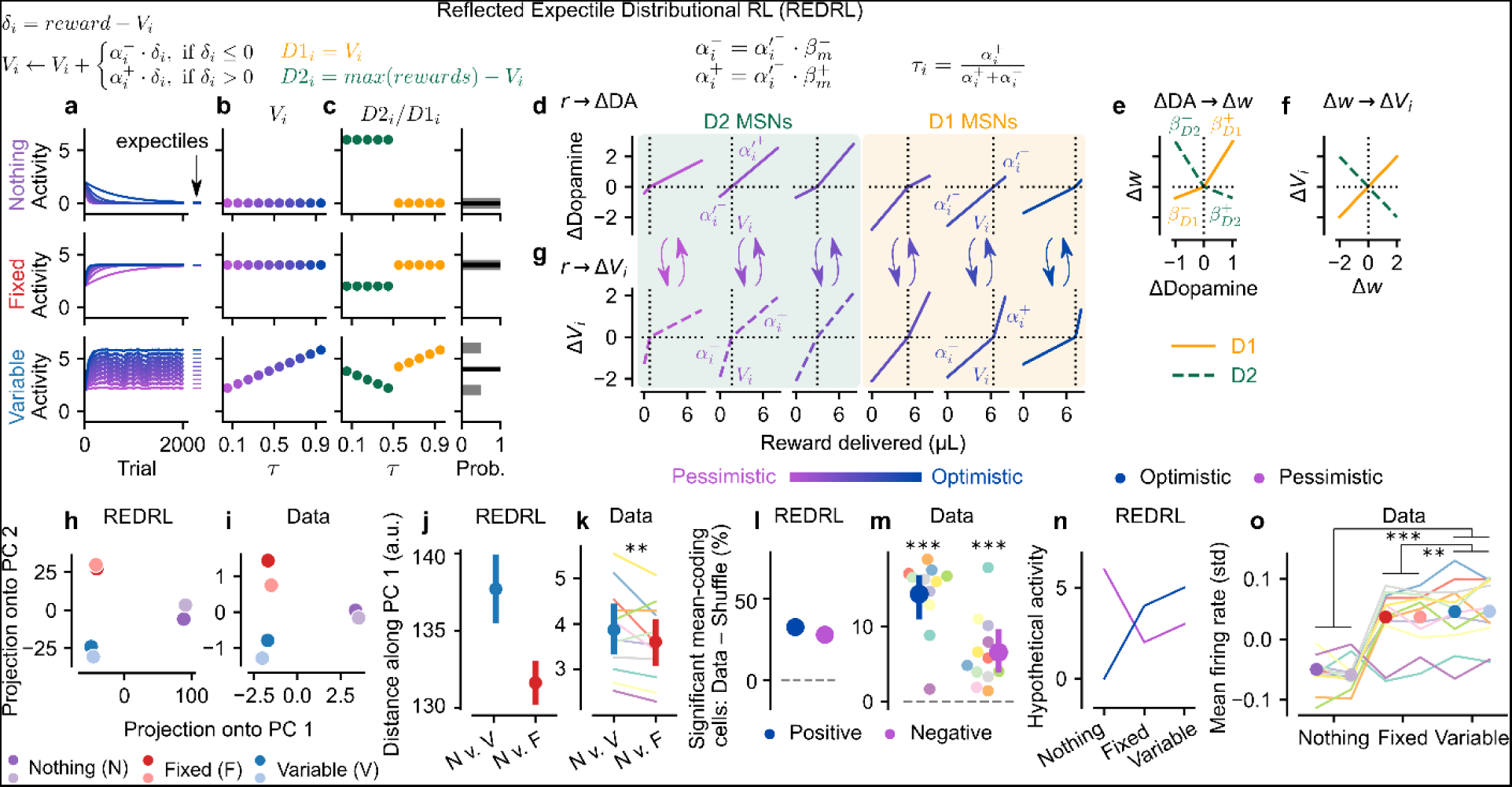
**| Reflected Expectile Distributional Reinforcement Learning (REDRL). a-c**, Algorithmic REDRL model. **a**, Over the course of training, value predictors (*V_i_*, here initialized to 2) converge to the expectiles of the associated reward distribution. **b**, Post-learning activity of the simulated value predictors, *V_i_*, as a function of their optimism level. The relative pessimism or optimism of each predictor is parameterized by τ, which can range from 0 to 1 (*x*-axis). **c**, *Left,* pessimistic (τ < 0.5) value predictors are identified with D2 MSNs (green), and their coding is flipped such that decreases in D2 activity correspond to increases in *V_i_*, and vice versa. Optimistic (τ > 0.5) predictors are directly proportional to D1 MSN activity (orange). *Right*, collectively, this striatal code characterizes the complete reward distribution via its expectiles. **d-g**, Implementation of REDRL within the mesolimbic circuit. **d**, Heterogeneity across dopamine neurons can be characterized using piecewise linear functions. Pessimistic neurons have high slopes in the negative domain (*α′_i_*^-^) and low slopes in the positive domain (*α′_i_*^+^), while the opposite is true for optimistic neurons. Over the course of learning, the zero-crossing point *V_i_* associated with each neuron will shift to equal the *τ_i_*-th expectile (vertical dotted line)^4^. **e**, D1 and D2 MSNs have asymmetric plasticity rules, potentiating more to increases and decreases in dopamine, respectively, relative to baseline (vertical dotted line)^39^. **f**, As a consequence, D1 activity is expected to correlate positively, and D2 activity negatively, with the corresponding value prediction^5,6^. To recover *V_i_*, we must subtract out the D2 activity, which could be accomplished for instance via its inhibitory projection to the ventral pallidum. **g**, The change in each value predictor is *ΔV_i_* = *α*′*_i_* ^−/+^.*β_m_*^−/+^.*δ* = *α_i_*^−/+.^*δ*, as demanded by the gradient descent-based update rule. The net result is that D1 MSNs are biased optimistically, and D2 pessimistically, relative to their dopamine input asymmetries, because their learning constants *β_m_*^-/+^ favor positive and negative prediction errors, respectively. With the plasticity rule shown, all D1 MSNs have *τ* > 0.5, and all D2 MSNs have *τ* < 0.5, justifying the division in **c**, though the precise distribution will depend on the specific plasticity rule and distribution of dopamine asymmetries. **h**, Two-dimensional PCA projection of converged value predictors, plus noise, for the REDRL model. Variable odors are further separated than Fixed from Nothing along PC 1 because after mean-centering, the patterns of Nothing and Variable activity are almost perfectly anticorrelated with one another, and the PC 1 loadings closely resemble Nothing activity itself. **i**, PCA projection of example session (same as Fig. 2b) shows a striking resemblance to the REDRL prediction in separating primarily Rewarded and Unrewarded odors along PC 1 and Fixed and Variable odors along PC 2. **j**, In addition, REDRL predicts that the distance between Nothing (N) and Variable (V) odors along PC 1 should be slightly greater than that between Nothing and Fixed (F). **k**, Striatal data are consistent with this prediction, with the distance along PC 1 significantly greater for Nothing vs. Variable than Nothing vs. Fixed odor pairs (*p* = 0.007). **l**, REDRL predicts that there should be substantial fractions of neurons that correlate either positively or negatively with mean value, corresponding to D1 and D2 MSNs. **m**, Significant populations of striatal neurons encode mean reward positively and negatively. Mean reward predicted on each trial was correlated with Late Trace activity. Then, for each neuron independently, we shuffled the odor-distribution mappings and re-computed the correlations. Each point denotes the per-mouse difference in fraction of significant cells (that is, cells with uncorrected *p* < 0.05) for the unshuffled and shuffled data, separately for cells that correlated positively or negatively with mean reward (Positive and Negative: *p* < 0.001, paired samples *t-* test comparing ordered and shuffled fractions across mice). **n**, REDRL predicts that Variable odors elicit higher population mean firing than Fixed odors, regardless of the optimism or pessimism of the underlying value predictor. **o**, Mean z-scored firing rates for each neuron, in addition to being higher for Rewarded than Unrewarded odors (*p* < 0.001), were also higher for Variable than for Fixed odors (*p* = 0.006), as assessed by an LME with neuron-level observations, averaged over trials, and mouse-level random effects.

Following observations of variance coding in other brain regions^87,89^, we identified variance-encoding neurons by linearly regressing single-neuron firing onto reward variance, after regressing out the effect of mean reward. Unlike these prior studies, however, we surprisingly found *fewer* striatal variance-encoding neurons than would be predicted from odor coding alone (Extended Data Fig. 3f-h). Furthermore, in contrast to codes in which neural activity is construed as representing samples from some probability distribution, across-trial Fano factors were the same across trial types with different variances^94,95^ (Extended Data Fig. 4a-d). We therefore adopted a different set of approaches to characterize distributional coding across the entire neural population.

**Fig. 4.**
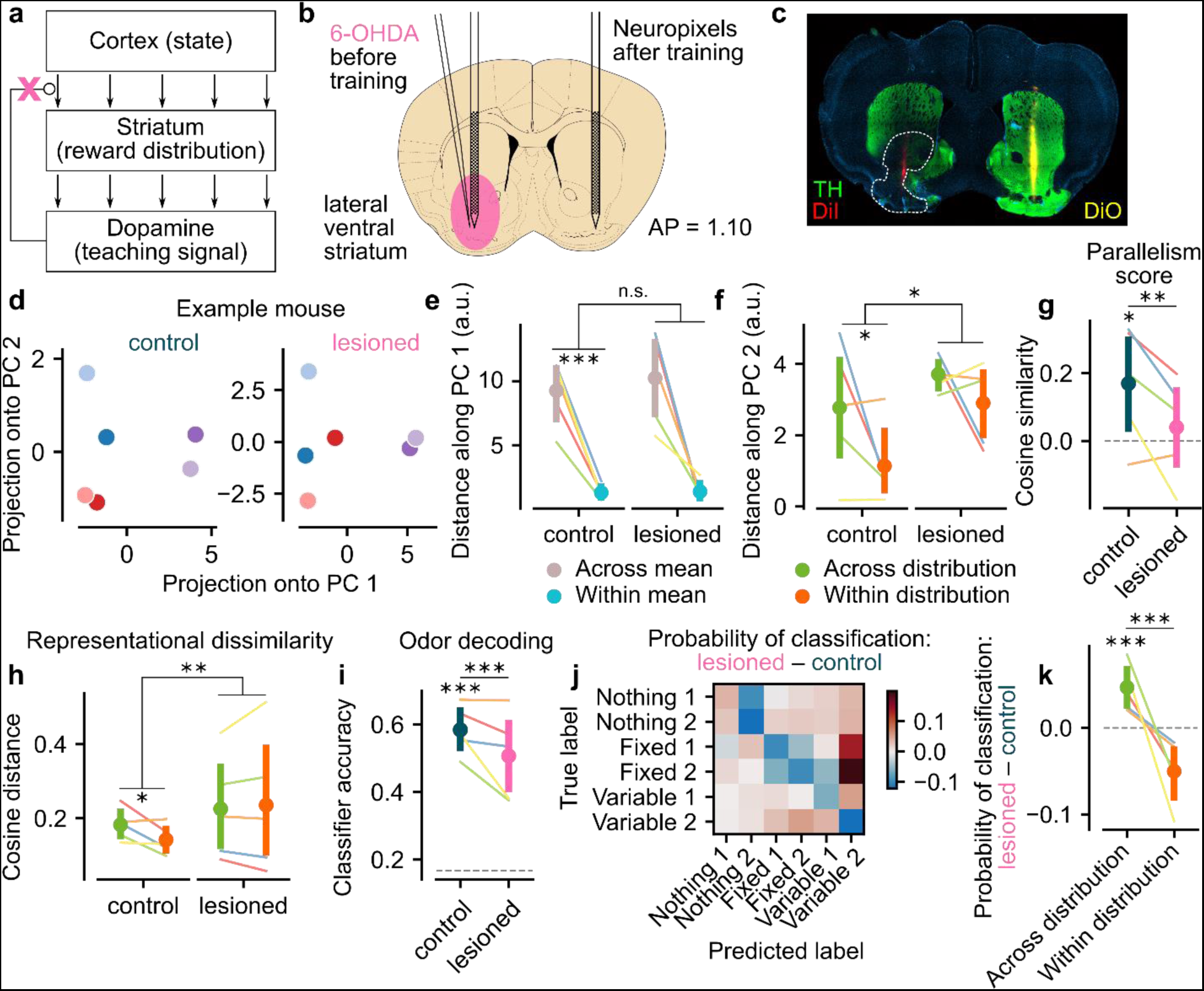
**| Dopamine is necessary for learning distributional representations. a**, Schematic illustration of the basal ganglia, showing how dopamine is hypothesized to act as a teaching signal to update corticostriatal synaptic weights. Therefore, dopamine lesions (pink “x”) are predicted to disrupt representations of the reward distribution in the striatum. **b**, Schematic illustration of dopamine lesion experiment. 6-OHDA was injected unilaterally into the lateral ventral striatum of naive mice to ablate dopamine neurons. After recovery and training, we recorded striatal activity in both the lesioned and control hemispheres. **c**, Histology from an example 6-OHDA animal showing Neuropixels probe tracks (red and yellow), dopamine axons (green), and lesion (white dashed line surrounding region of reduced TH staining). **d**, PCA projection of Late Trace activity from the control (*left*) and lesioned (*right*) hemispheres for an example mouse. **e**, Distance along PC 1, while significantly higher for across-mean than within-mean pairs (*p* < 0.001), does not differ between hemispheres (*p* = 0.676). For all panels of this figure, colored lines denote individual mice, averaged across pseudo-populations, and LMEs use these pseudo-populations as the individual observations with mouse-level random effects. **f**, By contrast, the difference in distance along PC 2 between across-and within-distribution pairs is significantly positive (*p* = 0.033) and greater for the control relative to the lesioned hemisphere (*p* = 0.026). **g**, Parallelism score is significantly positive (*p* = 0.029) and greater in the control relative to the lesioned hemisphere (*p* = 0.009). **h**, Similarly, the difference in representational dissimilarity between across-and within-distribution pairs is significantly positive (*p* = 0.036) and greater in the control relative to the lesioned hemisphere (*p* = 0.005). **i**, Six-way odor classification accuracy during the Odor period is above chance (*p* < 0.001) and higher for the control relative to the lesioned hemisphere (*p* < 0.001). **j**, Difference in odor classifier confusion matrices between the control and lesioned hemispheres. The probability of correct classification (main diagonal) decreases for nearly all trial types upon lesioning. **k**, The decrement in odor coding due to the lesion is mainly due to an increase in across-distribution, within-mean classification errors (the tendency in the lesioned hemisphere to predict Variable even when the true label was Fixed; *p* < 0.001) and a concomitant decrease in within-distribution classification (*p* < 0.001 for Across-vs. Within-distribution difference).

First, we projected each session’s trial type-averaged firing rates in the 1 s window before reward delivery (“Late Trace period”) onto the first and second PCs (accounting for 72.9 ± 2.4 and 10.0 ± 1.0% of the variance across trial types, respectively; mean ± s.e.m. across mice; Fig. 2b). We then measured the Euclidean distances in PC space along each dimension. As expected, trial types with different mean rewards segregated out along PC 1 (Fig. 2c). More surprisingly, though, Fixed and Variable odors separated out along PC 2, such that there was a greater distance between across-distribution odor pairs than within-distribution odor pairs (Fig. 2d).

Second, to determine whether we could observe the same trends in native firing rate space, we performed representational dissimilarity analysis (RDA) between the average population activity vector for each of the rewarded trial types (Fig. 2e). Once again, the distance between across-distribution pairs was greater, on average, than between within-distribution pairs (Fig. 2f). We observed the same effects in the classification performance of single-trial linear classifiers applied to pairs of rewarded trial types (Extended Data Fig. 5a-b) or applied to trial type groups that either respected or violated their distribution identities (Extended Data Fig. 5c-d). Distributional decoding was orthogonal to mean value coding (Extended Data Fig. 5e-h), stable over time (Extended Data Fig. 5i-k), and strongest in the more ventral and lateral parts of the striatum, particularly the lateral nucleus accumbens shell (lAcbSh; Extended Data Fig. 6). Lastly, an artificial neural network-based decoder trained on single pseudo-trial population activity successfully predicted complete reward distributions, even when its training and evaluation was restricted to trials with the same mean, and generalized to unseen odors (Extended Data Fig. 7).

**Fig. 5.**
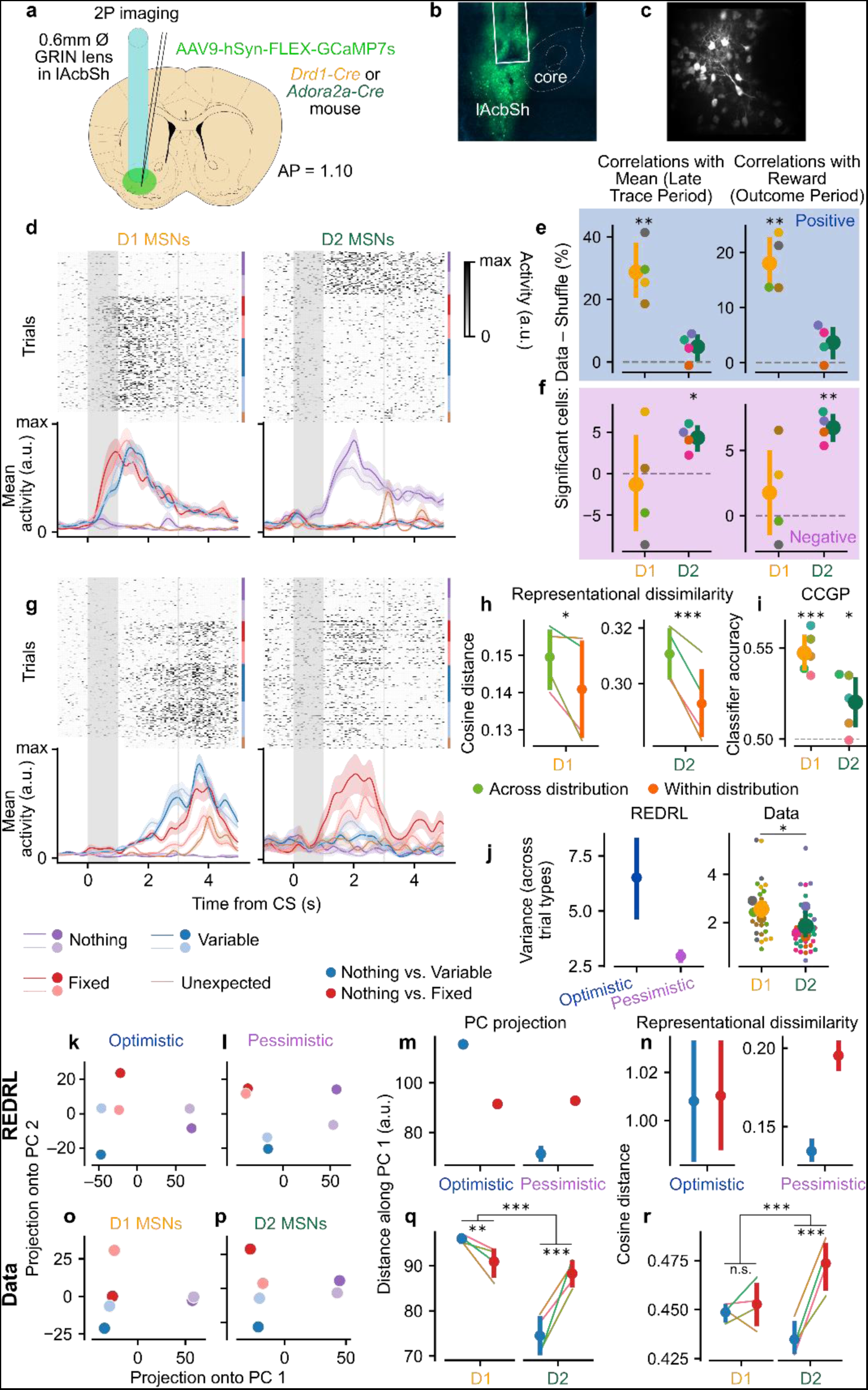
| Opponent contributions of D1 and D2 MSNs to distributional coding. a, Schematic illustration of two-photon calcium imaging experiment. We first injected a virus encoding the calcium indicator GCaMP7s and then implanted a GRIN lens in the lAcbSh in either *Drd1-Cre* or *Adora2a-Cre* mice, which drive Cre-dependent expression specifically in D1 and D2 MSNs, respectively. **b**, Example slice showing expression of GCaMP in the lAcbSh in a *Drd1-Cre* animal. **c**, Example FOV imaged through a GRIN lens in a *Drd1-Cre* animal. **d**, Deconvolved Ca^2+^ activity from an example D1 (*left*) and D2 (*right*) MSN. As in Fig. 2a, the top panel is a raster plot, normalized by maximum deconvolved activity, and the bottom panel shows average deconvolved activity ± s.e.m. across trials of each type. The D1 MSN responds most to Rewarded odors, while the D2 MSN responds most to Nothing odors. **e**, Quantification of average percentage of cells that correlate significantly positively with mean (*left*) or reward (*right*) during the Late Trace and Outcome periods, respectively, relative to the expectation from odor coding alone (shuffling odor-distribution mappings, horizontal dashed line). There are significantly more cells than expected by chance for D1 (paired samples *t*-test comparing ordered and shuffled fractions across mice, *p* = 0.009, 0.006 for mean and reward, respectively), but not D2 (*p* = 0.113, 0.107) MSNs. Thick lines show the mean ± 95% confidence interval across mice. **f**, Same as **e**, but for significant negative correlations. In this case, D2 (*p* = 0.013, 0.001) but not D1 (*p* = 0.736, 0.433) cells are significantly more common than expected by chance. **g**, Same as **d**, but showing an example D1 (*left*) and D2 (*right*) MSN that reliably discriminate Fixed and Variable odors. **h**, Cosine distance is significantly greater for across than within-distribution pairs for both D1 (*p* = 0.022) and D2 (*p* < 0.001) MSNs. For panels **h**, **i**, **q**, and **r** of this figure, individual replicates are pseudo-populations, split across trials and pooled across mice, hence there are no mouse-level random effects. Thick lines show the mean ± 95% confidence interval across pseudo-populations. **i**, CCGP is significantly above chance for both D1 (one-sample *t*-test relative to 0.5, *p* < 0.001) and D2 (*p* = 0.048) MSNs, demonstrating abstract encoding of variance in both populations. **j**, Variance across trial types, computed for the simulated REDRL predictors (*left*) and normalized neural data (*right*). Small dots are averages within sessions, medium dots are averages within mice, and large dots with error bars show averages ± 95% confidence intervals across mice (*p* = 0.017 for effect of genotype). **k-l**, Simulated REDRL value predictors were projected into two-dimensional PC space separately for optimistic (D1, **k**) or pessimistic (D2, **l**) value predictors. **m**, Quantification of Euclidean distance along PC 1 for the REDRL model. While optimistic predictors show the same trend as the complete code (Fig. 3j), pessimistic predictors swap the ordering between Fixed and Variable odors. Error bars denote 95% confidence intervals across odor pairs. **n**, Same as **m**, but using cosine distance in the full-dimensional space to quantify representational dissimilarity, again independently for optimistic and pessimistic predictors. **o-r**, Same as **k-n**, but showing data collected from D1 and D2 MSNs rather than simulated optimistic and pessimistic predictors, respectively. For both the distance along PC 1 (Nothing vs. Variable compared to Nothing vs. Fixed: *p* = 0.001 for D1, *p* < 0.001 for D2, *p* < 0.001 for the relative differences between D1 and D2) and the representational dissimilarity (*p* = 0.489 for D1, *p* < 0.001 for D2, *p* < 0.001 for the relative differences), striatal data closely match the theoretical predictions.

**Fig. 6.**
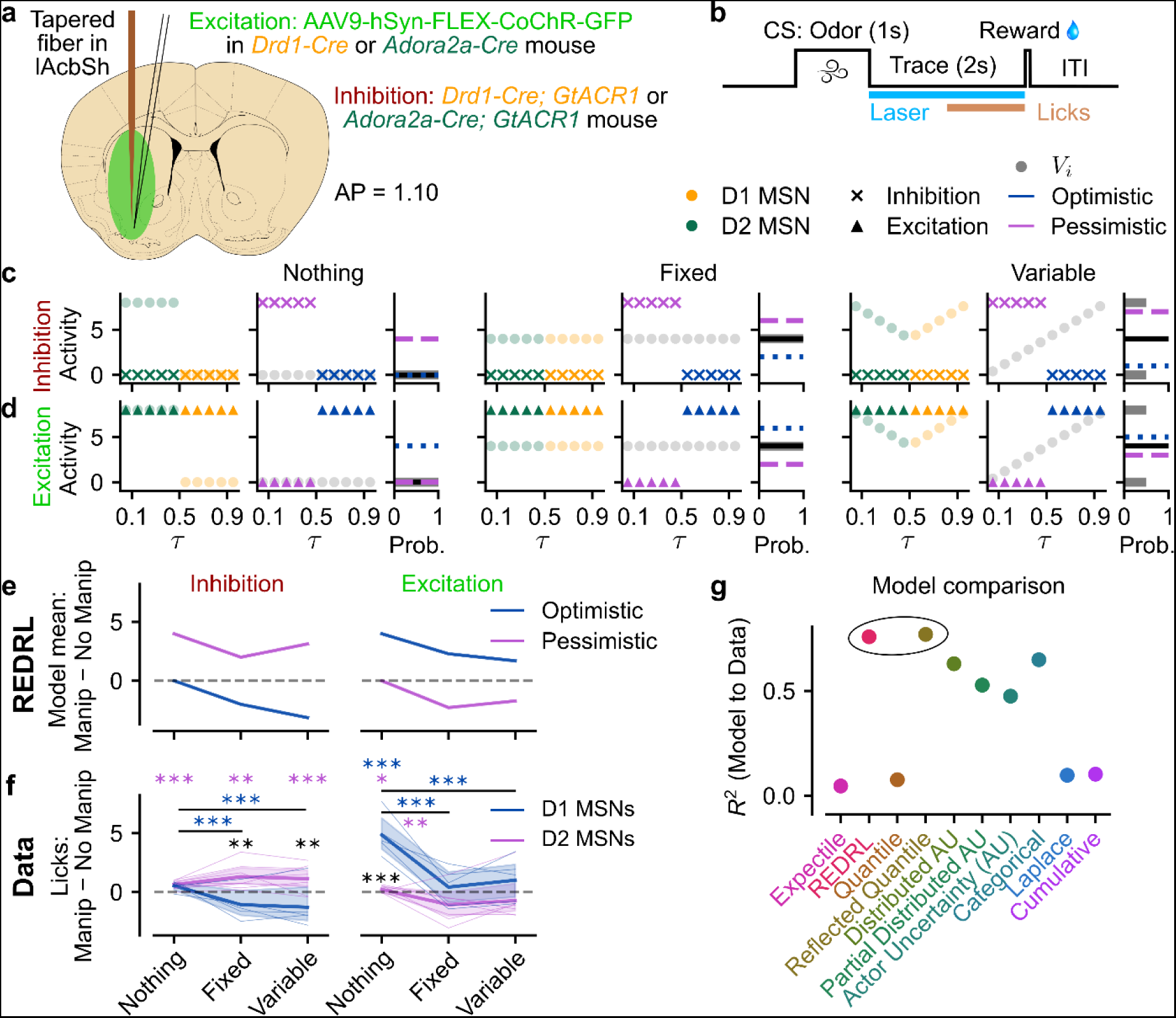
**| Causal contributions of D1 and D2 MSNs to REDRL. a**, Schematic illustration of optogenetics experiments. For excitation, a Cre-dependent virus containing the ultrasensitive excitatory opsin CoChR was injected into the lateral striatum at four separate depths. For inhibition, we used transgenic animals expressing the inhibitory opsin GtACR1, also in a Cre-dependent manner. Cre was delivered transgenically by way of *Drd1-Cre* or *Adora2a-Cre* mice, and a tapered fiber was implanted in the lAcbSh. **b**, The trial structure in these experiments was identical to the recording experiments, except that stimulation was delivered throughout the duration of the Trace period. Licking was again quantified during the Late Trace period, 1-0 s before the outcome, to avoid counting any artifactual licking around stimulation onset. The laser was pulsed for CoChR-based excitation and continuous for GtACR1-based inhibition. **c**, Approach for simulating the effects of optogenetic inhibition in the REDRL model. Within each group of panels (Nothing, Fixed, and Variable), the left column shows the predicted D1 (yellow) and D2 (green) activities for the No Manipulation (faded circles) and Manipulation (“x”s) conditions. Inhibition is simulated by clamping the relevant population to zero. The middle column portrays the resulting effect on the encoded value predictors, *V_i_*. In the REDRL model, optimistic (*τ* > 0.5; blue) and pessimistic (*τ* < 0.5; purple) predictors are identified with D1 and D2 MSNs, respectively. However, since the encoding of D2 MSNs is flipped, inhibition actually drives these *V_i_*’s positive relative to their baseline (grey). The right column illustrates the effect this change in *V_i_* has on the encoded mean (blue and purple horizontal dashed lines), relative to the unperturbed distribution (grey histogram, with mean shown in black). The ground-truth distributions shown reflect the versions used in the manipulation experiments, where the Variable condition consisted of equally probable 0 and 8 μL rewards. **d**, Same as **c**, but for optogenetic excitation (triangles) rather than inhibition. **e**, Summary of REDRL model predictions. Each point was computed as the difference in the implied mean between the Manipulation and No Manipulation conditions, computed separately for inhibition (*left*) and excitation (*right*). **f**, Difference in Late Trace period anticipatory licking between lAcbSh Manipulation and No Manipulation trials, computed within-session and then averaged across-session and within-mice (thin lines). Thick lines and shaded regions show the mean ± 95% confidence interval across mice. To emphasize the concordance with REDRL predictions, D1 and D2 manipulations are now colored blue and purple, respectively. Colored asterisks with horizontal lines denote significant differences in the effect of manipulation between trial types within the indicated genotype (D1 inhibition: *p* < 0.001 Nothing vs. Fixed, *p* < 0.001 Nothing vs. Variable; D1 excitation: *p* < 0.001 Nothing vs. Fixed, *p <* 0.001 Nothing vs. Variable; D2 excitation: *p* = 0.007, Nothing vs. Fixed). Colored asterisks over single trial types indicate significant differences relative to zero for that genotype (D2 inhibition: *p* < 0.001 Nothing, *p* = 0.002 Fixed, *p* < 0.001 Variable; D1 excitation: *p* < 0.001 Nothing; D2 excitation, *p* = 0.032 Nothing). Black asterisks over single trial types indicate significant differences between genotypes (inhibition: *p* = 0.001 Fixed, *p* = 0.005 Variable; excitation: *p* < 0.001 Nothing). **g**, Summary panel showing the mean coefficient of determination for each model, used to predict the average difference in licking for each trial type without any intercept term.

## Variance is encoded abstractly

The preceding analyses show that the neural activities evoked by odors identifying the same distribution are more similar to one another than to those evoked by odors identifying distributions with the same mean but different variances. Let us now ask about the *relationship* between Fixed and Variable odor representations. More specifically, is variance represented in an “abstract format” — i.e., in a way that supports generalization to unseen situations^96^? To find out, we adapted two previously-defined metrics^96^ to our task: parallelism score and cross-condition generalization performance (CCGP). Both ask, in different ways, whether there is a consistent direction in firing rate space that distinguishes low and high-variance cues (see Methods).

The parallelism score is simply the average cosine similarity between the two difference vectors pointing from Variable to Fixed population activity, one for each odor identifying the respective distribution (Fig. 2g). Across sessions and mice, these difference vectors were significantly more aligned than would be expected by chance (Fig. 2h). Similarly, a decoder trained on one Fixed vs. Variable dichotomy and then tested on the held-out dichotomy achieved above-chance CCGP, averaged across all four possible dichotomies (Fig. 2i-j). These analyses show that variance is not just encoded arbitrarily, but in an abstract format.

## Using striatal opponency to implement distributional RL

How might such an abstract representation be acquired? While there exist multiple theories for how the brain might learn (factorized, that is, abstract) reward distributions^71,72,83^, EDRL^76^ is especially promising because it requires only minimal modifications to existing, empirically tested models of the basal ganglia^4^. EDRL proposes not just a single value predictor but an entire family of predictors *V_i_*, which learn at different rates, *α_i_^+^* and *α_i_*^-^, for positive and negative RPEs, respectively (Fig. 3a). “Optimistic” predictors have relatively high *α_i_^+^* and will converge to values above the distribution mean, while the opposite is true of “pessimistic” predictors. Each predictor converges to a so-called “expectile” of the reward distribution, parameterized by 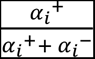 between 0 and 1. Expectiles generalize the mean (*τ* = 0.5) just as quantiles generalize the median, and collectively, they characterize the complete reward distribution^97^ (Fig. 3b; see Methods).

While EDRL has some appealing properties, it ignores the molecular and cellular diversity within the striatum, most notably the presence of D1 and D2 MSNs^38^. As an extension, we propose reflected EDRL (REDRL) — so called because D2 MSN activity is simply the negative of the corresponding value predictor, plus a constant offset to ensure non-negative activities (Fig. 3c). This simple modification does not merely lend EDRL additional biological plausibility; rather, it demonstrates how the particular anatomy of the striatum can benefit distributional RL computations while explaining a host of data regarding activity in the striatum and opponency between D1 and D2 MSNs.

To implement REDRL in the striatum, we first require structured heterogeneity in dopamine inputs, which can be modeled as piecewise linear response functions to reward size^4^ (Fig. 3d). Since RPE is defined as actual minus predicted reward, the reward amount which elicits no change in dopamine firing relative to baseline — the so-called “zero-crossing point”^4^ — is equivalent to the learned value prediction for that neuron. Pessimistic dopamine neurons have steeper slopes for rewards below their associated value prediction (*α′_i_*^-^) and shallower slopes above it (*α′_i_*^+^), reflecting relatively low learning rates from positive RPEs. The converse is true of optimistic dopamine neurons. Second, these diverse dopamine responses combine with opponent plasticity rules in D1 and D2 MSNs, with D1 MSNs increasing synaptic weights more from positive RPEs (*β_m_ ^+^*) and D2 MSNs increasing synaptic weights more from negative RPEs^22,39,40^ (*β_m_* ^-^; Fig. 3e). Importantly, while asymmetric, these synaptic weight updates are not fully dichotomous; D1 and D2 MSNs still learn slightly from dopamine changes in their non-preferred directions^66^, in line with the shallower but nonzero slope of D1 and D2 receptor occupancy curves at baseline dopamine concentrations^66,98,99^.

By composing these two functions, we get the complete REDRL model. The *opponency* of the plasticity rule gives rise to opponent directions of value coding (Fig. 3f), with D1^5,6,100,101^ and D2 MSNs^5,6,102^ primarily correlating positively and negatively, respectively, with value and reward. Meanwhile, its *asymmetric* nature has the effect of extremizing value predictors — D1 MSNs are more optimistic, and D2 MSNs more pessimistic, than their individual dopamine inputs would create on their own (Fig. 3g) — setting up a global bias for D1 and D2 MSNs to encode the right and left tails of the value distribution, respectively, and shifting the zero-crossing points of the coupled dopamine neurons up or down accordingly (Fig. 3d-g). Notably, this also predicts that D1 MSNs will acquire positive associations faster than D2 MSNs, while D2 MSNs may be preferentially involved in later discrimination or extinction^39,40,103^ (Fig. 3a).

Armed with such a model, we can ask whether the population activity predicted by REDRL mirrors that observed in our striatal data. Strikingly, the top two PCs of the model predictors closely resemble the projection of the data using principal component analysis (PCA; Fig. 3h-i). Moreover, REDRL gives rise to a new prediction: Variable odors should be more distant from Nothing odors along PC 1 than Fixed odors, a prediction that due to PCA’s mean-centering is independent of the D2 offset, and that we confirmed to hold true in our data (Fig. 3j-k). Secondly, REDRL predicts the existence of substantial populations of neurons that correlate either positively (D1) or negatively (D2) with expected reward across trials (Fig. 3l). We again found this to be the case in our data (Fig. 3m), with a slight bias toward positive correlations, perhaps reflecting the preponderance of D1 over D2 MSNs in the striatum^104-106^. Lastly, REDRL predicts that the average firing rate should be slightly higher for Variable than for Fixed odors on average, which we also observed (Fig. 3n-o). While other distributional RL formulations predicted some of these effects, only REDRL and its close cousin, Reflected Quantile Distributional RL (Extended Data Fig. 8a-m) predicted all of them. Thus, REDRL provides a mechanistic account of distributional reinforcement learning which quantitatively matches the structure of striatal representations.

## Dopamine is necessary for distributional RL

If striatal representations are updated incrementally by dopamine RPEs as predicted by REDRL, then eliminating dopamine prior to learning should disrupt these distributional representations (Fig. 4a). To test this hypothesis, we injected the neurotoxin 6-hydroxydopamine (6-OHDA) unilaterally into the lateral ventral striatum in naive mice, which resulted in local lesions of dopamine neurons projecting to the injection site (Fig. 4b-c; Extended Data Fig. 9a). After recovery, we trained the animals on the same task and then recorded neurons in both the control and lesioned hemisphere (*N* = 5 mice, *n* = 20 sessions, 2,283 neurons from control; 19 sessions, 2,596 neurons from lesion). Unilateral lesions modestly impaired our ability to distinguish Rewarded and Unrewarded odors based on behavioral predictors, but animals nonetheless learned the task (Extended Data Fig. 9b-c).

Projecting striatal activity from each hemisphere independently into PC space suggested that Fixed and Variable distributions were less well-separated in the lesioned hemisphere relative to the control hemisphere (Fig. 4d). Indeed, when we quantified distances as before, we found Nothing and Rewarded odors to be equally well separated along PC 1 for both hemispheres (Fig. 4e), but less-well separated along PC 2 in the lesioned hemisphere, with an associated smaller difference in distances between across-distribution and within-distribution pairs (Fig. 4f). Analogous effects were seen for parallelism score (Fig. 4g) and representational dissimilarity (Fig. 4h), with stronger (and abstract) variance coding in the control relative to the lesioned hemisphere. The persistence of mean value coding in the lesioned hemisphere may reflect the inability of unilateral 6-OHDA to kill all dopamine neurons within the targeted hemisphere, the interhemispheric broadcasting of mean value information once it reaches cortex^31-37^, or, more radically, the dispensability of dopamine for learning about mean value entirely.

In addition to supporting our mechanistic REDRL model, the selective disruption of variance coding by 6-OHDA gives us an experimental tool with which to probe the function of distributional RL in the brain. When paired with deep neural networks, distributional RL is thought to boost system performance mainly by improving state representations^1,4,78^. Due to multiplexing of odor-specific representations alongside distribution information within the striatum (Extended Data Fig. 3), it is possible to ask whether dopamine lesions also impair striatal stimulus representations. We used multinomial logistic regression to decode odor identity from neural activity during the 1 s window following odor onset. While we could decode odor identity well above chance for both hemispheres, decoding performance was significantly higher in the control than the lesioned hemisphere (Fig. 4i). The lesion caused a drop in decoding performance across nearly all trial types, with the main driver being an increased confusion between Fixed and Variable odors (Fig. 4j-k). These results are consistent with distributional RL playing a similar role in shaping the representation of sensory inputs in artificial neural networks and biological brains.

## Opponent contributions of D1 and D2 MSNs to REDRL

To dissect the distinct contributions of D1 and D2 MSNs predicted by REDRL, we turned to two-photon calcium imaging through implanted gradient refractive index (GRIN) lenses (Fig. 5a). We injected *AAV9-hSyn-FLEX-jGCaMP7s* virus^107^ into the lAcbSh of *Drd1-Cre* (*N* = 4 mice, *n* = 27 sessions, 945 neurons) or *Adora2a-Cre* (*N* = 4 mice, *n* = 38 sessions, 1,106 neurons) transgenic mice^108^, which drive expression in D1 and D2 MSNs, respectively (Fig. 5b). Using this method, we were able to image up to ∼50 neurons simultaneously per field of view (31.6 ± 17.4, mean ± s.d. across sessions; Fig. 5c).

We observed different patterns of deconvolved Ca^2+^ activity across D1 and D2 populations^100-102^ (Extended Data Fig. 10a-b). Many D1 MSNs were activated more to Rewarded than to Unrewarded odors and outcomes, while the reverse was true, albeit less strongly, in D2 MSNs (Fig. 5d-f). In addition to correlations across trials during the Late Trace period, we also investigated differences between the Late Trace and Baseline periods, because REDRL predicts increases in value predictions after Rewarded odor onset. Consistent with our model, significant fractions of D1 and D2 MSNs increased and decreased their activities, respectively, more on Rewarded than Unrewarded trials, although the pattern in D2 MSNs was again more heterogeneous than in D1 MSNs (Extended Data Fig. 10c).

Intriguingly, we also found neurons which, like those we recorded using electrophysiology, reliably distinguished between Fixed and Variable odors during the Late Trace period (Fig. 5g). To test whether these trends were systematic, we performed the same analyses (RDA, CCGP and PCA) separately on D1 and D2 MSNs, while pooling across all sessions and all mice to compensate for the lower cell counts and higher variability of Ca^2+^ signals. Consistently across disjoint subsets of trials in both D1 and D2 MSNs, across-distribution pairs were represented more dissimilarly than within-distribution pairs (Fig. 5h), and variance was encoded in an abstract format (Fig. 5i).

REDRL not only predicts that distributional coding should be present in both D1 and D2 MSNs independently but also specifies the ways in which this coding should differ. For one, this particular set of distributions should elicit higher variance across trial types for optimistic than for pessimistic reward predictors on average — which is also true in the two-photon data for D1 and D2 MSNs, respectively (Fig. 5j). More impressively, when projecting optimistic (*τ* > 0.5) and pessimistic (*τ* < 0.5) predictors into 2D PC space separately, we found that optimistic predictors exhibited the same trend as the full complement of value predictors, with Variable and Nothing odors further separated along PC 1 than Fixed and Nothing odors (Fig. 5k). However, pessimistic predictors actually showed the opposite trend, with Variable and Nothing odors closer together along PC 1 than Fixed and Nothing (Fig. 5l-m). Analogously, representational dissimilarity was less for Variable and Nothing odors than Fixed and Nothing odors specifically for pessimistic predictors; optimistic predictors did not differ (Fig. 5n). PCA projections (Fig. 5o-q) and RDA (Fig. 5r) for D1 and D2 MSNs mirrored these predictions precisely, revealing a subtle distinction in distributional coding across MSN subtypes and confirming a novel prediction of REDRL.

## Perturbing REDRL with optogenetics

As a final test of REDRL, we sought to independently manipulate D1 and D2 MSNs while mice performed a similar classical conditioning task. To do so, we expressed either the excitatory opsin CoChR^109^ (*N* = 12 mice, *n* = 96 sessions) or the inhibitory opsin GtACR1^110,111^ (*N* = 13 mice, *n* = 92 sessions) in D1 or D2 MSNs and implanted an optical fiber in lAcbSh^112^ (Fig. 6a). We then manipulated these neurons during the 2 s Trace Period after odor offset and quantified licking during the last 1 s of this Trace Period, just prior to reward delivery (Fig. 6b).

To identify the REDRL model predictions for these manipulations, we clamped the simulated values of inhibited and excited predictors respectively at 0 and 8 μL, the maximum reward size we delivered in these experiments. We performed these simulated manipulations separately in optimistic and in pessimistic neurons while letting the non-manipulated predictors retain their original values. We then computed the animal’s predicted value estimate as the mean across both optimistic and pessimistic predictors (Fig. 6c-d). For comparison, we performed similar manipulations on other distributional code types (Extended Data Fig. 11). We then took the difference between the models’ estimated values in Manipulation vs. No Manipulation trials for each trial type (Fig. 6e; Extended Data Fig. 12) and compared it to the difference in anticipatory licking. REDRL not only captured the main effects of “go” and “no-go” pathways^113^ but also predicted precise patterns of licking across trial types, even for the same type of manipulation (Fig. 6f). This could not be explained simply by ceiling effects, as the increase in licking was sometimes greater for Rewarded than Unrewarded odors, as in the case of D2 inhibition. Quantitative comparison between the data and various models confirmed that REDRL (and the highly similar Reflected Quantile code) best fit the licking data (Fig. 6g).

## Discussion

Here we have combined large-scale electrophysiology with cell-type specific recordings and manipulations to develop the REDRL model of the basal ganglia. This model maintains the normative algorithmic advantages of distributional RL^1^ while lending itself to a biological implementation that is consistent with the observed structure of dopamine population activity^4^ and dopamine-mediated plasticity rules^22,39,40^. The most notable feature of REDRL is the distinct role played by D1 and D2 MSNs, which specialize in the right and left tails of the reward distribution, respectively. This bifurcated layout resembles other neural systems, such as ON/OFF pathways in vision, and likely has similar benefits, such as efficient coding^114^, reduced metabolic cost^115^, and flexibility^116^. For example, certain computations, such as expected value estimation, would benefit from combining information from D1 and D2 MSNs, but others, such as risk-sensitive behavior, might depend on one tail or the other, and thus primarily require information from a single population. Furthermore, this architecture simplifies the problem of connectivity: anatomically and/or genetically-defined subsets of dopamine neurons^117,118^ could form independent closed loops with D2 (via ventral pallidum) and D1 MSNs, thereby helping to keep separate pessimistic and optimistic RPE channels. These predictions should form the basis of future anatomical investigations into the mesolimbic dopamine circuitry, as well as theories of alternative architectures that might obviate this need^119^, which is shared by EDRL.

At the level of the striatum, REDRL helps unify previous approaches to understanding D1 and D2 MSNs within a single, normative framework. While there have been previous hints that D1 and D2 MSNs are oppositely modulated by dopamine^22,39,40^ and oppositely correlated with reward and expected value^5,6,100-102^, this has generally been attributed to go/no-go or approach/avoid pathways and modeled using a single value predictor^3,66,113,120,121^. Here, we show how, far from being a bug or redundancy in the RL architecture, such diversity could actually be a feature, biasing convergence to optimistic or pessimistic value predictors. More speculatively, it could explain why D1 and D2 MSNs often act in an opponent fashion without being inverses of each other^41-45^. The tendency for both pathways to activate prior to movement onset, for example, would be predicted if such transition points coincide with increases in the predicted variance of rewards (and thus the density on both the left and right tails).

REDRL also lends a new perspective to the coding of uncertainty in the brain. Typical treatments of this topic focus on *perceptual* uncertainty, where the observer’s role is to infer the distribution of world states consistent with a pattern of neural activity^70^. While the problem is generally formulated as one of Bayesian inference^67^, the associated uncertainty is frequently attributed to noisy inputs rather than ones that are genuinely ambiguous (as in the case of the Necker cube^122^). In RL settings, in contrast, uncertainty generally arises from a combination of state ambiguity, insufficient exploration, and intrinsic stochasticity^123^, all of which complicate the problem of learning from limited experience. Distributional RL excels in partitioning out this intrinsic uncertainty from other sources, potentially allowing for improvements in state representation^77,78^, exploration^79-82^, value estimation^124^, model-based learning^125^, off-policy learning^126^, and risk sensitivity^127-130^.

Many questions remain as to how the brain transforms high-dimensional reward distributions into a single choice, but it is tempting to speculate that this process corresponds to the dimensionality reduction that takes place throughout the various nuclei of the basal ganglia^131^, ultimately collapsing onto a unitary value estimate in the mediodorsal thalamus that defines the choice axis. Notably, such a “distributional critic” — centered here in the lAcbSh, a region which receives RPE-like mesolimbic dopamine input^46-50^ — could integrate seamlessly into a broader RL framework^132-136^, with the dorsal striatum likely playing the role of the “actor” and choosing actions in continuous, high-dimensional spaces^137^. Modifications of the encoded reward distribution, such as by dopaminergic drugs^59,60^, or of the downstream basal ganglia circuit, could then bias risky choice on rapid or developmental timescales^61,138,139^. Various psychopathologies — such as depression, in which patients learn more from losses than gains^140,141^, or addiction, in which patients systematically overweight the right tail of the reward distribution^142^ — could similarly stem from the dysfunction of this core distributional RL circuitry. Thus, REDRL can serve as a bridge between reinforcement learning, behavioral economics, computational psychiatry, and systems neuroscience, demonstrating how the circuit logic of the striatum can combine with vector-valued dopamine signals to realize the computational benefits of distributional RL.

## Methods

### Experimental Procedures

#### Mice

A total of 46 adult C57BL/6J (Jackson Laboratory) male and female mice were used in these experiments. Twelve wildtype animals (6 M, 6 F) were used for Neuropixels recordings, of which five (2 M, 3 F) were also included in unilateral 6-OHDA experiments. For two-photon imaging, four *Drd1-Cre* (B6.FVB(Cg)-Tg(Drd1-cre)EY262Gsat/Mmucd, RRID:MMRRC_030989-UCD; 3 M, 1 F) and four *Adora2a-Cre* (B6.FVB(Cg)-Tg(Adora2a-cre)KG139Gsat/Mmucd, RRID:MMRRC_036158-UCD; 1 M, 3 F) mice were used^108,143,144^. For optogenetic excitation, we used five *Drd1-Cre* (2 M, 3 F) and seven *Adora2a-Cre* (3 M, 4 F) animals. For optogenetic inhibition, we crossed these lines with a Cre-dependent *GtACR1* reporter mouse^111^ (R26-CAG-LNL-GtACR1-ts-FRed-Kv2.1, RRID:IMSR_JAX:033089). Five *Drd1-Cre;GtACR1* (2 M, 3 F) and eight *Adora2a-Cre;GtACR1* (4 M, 4 F) mice were used. All transgenic mice used for experiments were backcrossed with C57BL/6J and heterozygous for the relevant allele(s).

Animals were housed on a 12 hr dark/12 hr light cycle and performed the task at the same time each day (± 1 hour), during the dark period. Ambient temperature was kept at 75 ± 5°F, and humidity was kept below 50%. Animals were group-housed (2-5 animals/cage) until surgery, then individually housed throughout training and testing. All procedures were performed in accordance with the National Institutes of Health Guide for the Care and Use of Laboratory Animals and approved by the Harvard Institutional Animal Care and Use Committee (IACUC).

#### Surgeries

All surgeries were performed under aseptic conditions. Mice (> 8 weeks old) were anesthetized with isoflurane (3.5% induction, followed by 1-2% maintenance at 1 L/min), and local anesthetic (lidocaine, 2%) was administered subcutaneously at the incision site. Analgesia (buprenorphine for pre-operative treatment, 0.1 mg/kg, intraperitoneal (i.p.); ketoprofen for post-operative treatment, 5 mg/kg i.p.) was administered for two days after surgery. After leveling, cleaning, and drying the skull, we affixed a custom-made titanium head plate to the skull with adhesive cement^20^ (C&B Metabond, Parkell).

For all injections, the solution (6-OHDA or virus) was backfilled into a pulled glass pipette (Drummond, 5-000-1001-X), followed by mineral oil and a plunger. A small craniotomy (< 1 mm diameter) was made using a dental drill, and then the pipette assembly was mounted on the stereotaxic holder, lowered to the desired coordinate, and injected slowly (∼100 nL/min) to minimize damage to the surrounding tissue (Narishige, MO-10). After each injection, we waited at least 10 minutes to allow the solution to diffuse away from the pipette tip before slowly going up to the next coordinate or retracting the pipette from the brain. Target coordinates (in mm) for the lAcbSh were the same across experiments: AP 1.1 from bregma, ML 1.7, and DV 4.2 from the pial surface.

#### 6-OHDA procedure

To unilaterally ablate dopamine neurons projecting to lateral ventral striatum, we followed an existing protocol^52,145^. The following solution was injected (i.p.) into animals at 10 mg/kg prior to surgery:

● 14.25 mg desipramine (Sigma-Aldrich, D3900-1G)
● 3.1 mg pargyline (Sigma-Aldrich, P8013-500MG)
● 5 mL distilled water

Most animals (weighing ∼25 g) received ∼250 μL of this solution, which was given to prevent dopamine uptake in noradrenaline neurons and to increase the selectivity of update by dopamine neurons. We additionally prepared a solution of 10 mg/mL 6-hydroxydopamine (6-OHDA; Sigma-Aldrich, H116-5MG) and 0.2% ascorbic acid in saline (0.9% NaCL; Sigma-Aldrich, PHR1008-2G). The ascorbic acid in this solution helps prevent 6-OHDA from breaking down. The control hemisphere was either injected with vehicle ascorbic acid solution or uninjected; we observed no differences between these groups and so combined them. To further prevent 6-OHDA from breaking down, we kept the solution on ice, wrapped in aluminum foil, and used it within three hours of preparation. If the solution turned brown during this time (indicating that 6-OHDA had broken down), it was discarded and a fresh solution was made. 225 nL 6-OHDA (or vehicle) was injected unilaterally into lAcbSh.

Surgeries occurred at least 1 week before the start of behavioral training. We lesioned nine animals and included control hemisphere data for all of them in the main dataset. However, four of these animals either died before we could record from the lesioned hemisphere or were not correctly targeted for the lesion and/or recording, and so were excluded from the lesion dataset.

#### Viruses

To express constructs specifically in D1 or D2 MSNs, we injected viruses into *Drd1-Cre* and *Adora2a-Cre* mice. For imaging experiments, we unilaterally injected 450 nL AAV9-hSyn-flex-GCaMP7s (≥ 1×10¹³ vg/mL, Addgene)^107^ into lAcbSh. For optogenetic activation experiments, we bilaterally injected AAV9-hSyn-flex-CoChR-GFP (5.1 x 10^12^ vg/mL, UNC Vector Core, NC)^109^ at AP 1.1, ML ±1.7 in 300 nL increments at four separate depths below the pial surface: 4.2, 3.4, 2.6, and 1.8.

#### GRIN lens and fiber implantations

Prior to GRIN lens surgery we injected animals i.p. with 50 μL dexamethasone (2 mg/mL; Vedco) to reduce inflammation. Before virus injection, a needle was mounted on the stereotaxic holder, connected to light suction, and lowered to 3.4 mm below the pial surface to gently aspirate away the overlying brain tissue. After virus injection, a singlet GRIN lens (0.5 NA, 0.6 mm diameter, 7.3 mm length, 0 - 200 µm WD, 3/2 pitch, Inscopix, 1050-004597) was mounted onto a stereotaxic cannula holder (Doric) and then slowly lowered over at least 30 minutes to its target depth, 200 μm above the injection site and 3.8 mm below the pial surface. Metabond was used to secure the GRIN lens on all sides and allowed to dry completely before removing the cannula holder and covering everything with another layer of Metabond mixed with charcoal powder to block out light. Lastly, a plastic cap was attached with Kwik-Cast (World Precision Instruments) to protect the lens from damage.

For optogenetic manipulation, we bilaterally implanted tapered fibers (0.66 NA, 200 μm diameter, 3 mm emitting length, 5 mm implant length; Optogenix) in the lAcbSh after virus injection, at a depth of 4 mm. Each fiber was secured using Metabond and then protected with a fitted cap.

#### Behavior setup and tasks

Behavioral events were controlled (and licking was monitored) using custom-written software in MATLAB (Mathworks, Natick, MA) and the Bpod library (Sanworks, Rochester, NY) interfacing with the Bpod state machine (Sanworks, 1024 and 1027), valve module (Sanworks, 1015), and port interface board (Sanworks, 1020)/water valve (Lee Company, LHDA1233115H) assembly. Odors were delivered using a custom olfactometer^146^, which directed air through one of eight solenoid valves (Lee Company, LHDA1221111H) mounted on a manifold (Lee Company, LFMX0510528B). Each odor was dissolved in mineral oil at 10% dilution, and 30 μL of diluted odor solution was applied to a syringe filter (2.7 μm pore, 13 mm diameter; Whatman, 6823-1327). Wall air was passed through a hydrocarbon filter (Agilent Technologies, HT200-4) and split into a 100 mL/min odor stream and 900 mL/min carrier stream using analog flowmeters (Cole-Parmer, MFLX32460-40 and MFLX32460-42), which were recombined at the odor manifold before being delivered to the animal’s nose. Licking was monitored using an infrared emitter-photodiode pair positioned just in front of the plastic lick spout, positioned at the animal’s mouth.

Animals used for Neuropixels recording and 2-photon imaging were conditioned with six different neutral odors, chosen at random from these seven: isoamyl acetate, *p*-cymene, ethyl butyrate, (*S*)-(+)-carvone, (±)-citronellal, α-ionone, and L-fenchone. Optogenetic manipulation animals used only the first three. In all experiments, the mapping between physical odor and conceptual trial type was randomized across mice. Each trial began with a 1 s odor presentation, followed by 2 s trace period and then reward delivery. There was a minimum of 4.6 s before the next trial (4.1 s for optogenetic manipulation animals), plus a variable ITI drawn from a truncated exponential distribution with a mean of 2 s, minimum of 0.1 s, and maximum of 10 s. For 2-photon imaging experiments, this was extended to a mean of 10.5 s, minimum of 6.5 s, and maximum of 18.5 s to account for the slower kinetics of the calcium indicator relative to electrophysiology.

The recording task consisted of three different reward distributions, Nothing, Fixed, and Variable (Fig. 1b). Each distribution was then paired with two unique odors, for a total of six odors. The distributions were as follows:

● Nothing: 100% chance of 0 μL water
● Fixed: 100% chance of 4 μL water
● Variable: 50% chance of 2 μL water; 50% chance of 6 μL water

The task used for optogenetic manipulation was simplified in two ways. First, we used only one odor per distribution, for a total of three odors. Second, we modified the Variable distribution to be 50/50% between 0 and 8 μL, because our model predicted that increasing the variance would lead to a greater behavioral difference between Fixed and Variable odors.

#### Behavior training

Water restriction began no earlier than 5 days after recovery from surgery. Animals’ condition was monitored daily to ensure that mice did not dip below 85% of their free-drinking body weight, including supplementing with additional water after the task to bring their total daily intake to ∼1.2 mL. Over the course of three successive habituation days, mice were (1) handled gently for several minutes in their home cage, (2) permitted to freely roam around the platform in the behavior rig to collect water and then (3) head-fixed while receiving frequent (inter-reward interval 4-5 s) 6 μL water rewards.

The optogenetic manipulation task proceeded in only one phase, with up to 110 Nothing, 110 Fixed, and 114 Variable trials, randomly interleaved. By contrast, training for the recording task took place in three phases, each with a maximum of 300 trials.

● In Phase 1, mice experienced both Nothing odors and both Fixed odors with equal probabilities
● In Phase 2, mice experienced all six odors, but with the Variable odors 5.5x more frequent than the others
● In Phase 3, mice experienced all six odors at the final ratio of 4:4:7 (Nothing:Fixed:Variable), to increase the statistical power for analyzing responses to different reward sizes

On recording days, animals experienced a maximum of 20 additional Unexpected reward trials, in which 4 μL of water was delivered without being preceded by an odor cue. All trials were randomly interleaved in all phases.

For both tasks, animals completed at least 150 trials per day, and almost always more than 250. The experiment might be terminated early by the experimenter if the animals stopped licking in anticipation (or consumption) of the rewards due to satiety. A behavior session was considered “significant” if the lick rate during the last half second prior to reward delivery was significantly different between Rewarded (Fixed and Variable) and Unrewarded (Nothing) odors (Mann-Whitney U test, α = 0.05) and the effect size was at least 0.75 licks/s. Animals were advanced to the next phase, or to habituation for recording/manipulation, after at least two consecutive days with significant behavior. On recording/manipulation days, only significant behavior sessions were included for neural or behavioral analysis.

#### Neuropixels recordings

The day before recording, animals were habituated to the recording setup by covering their heads with a plastic sheet to block their view of the probe and manipulator. We then turned on the lamp, ran the brushed motor controller (Thorlabs, KDC101 and Z825B) up and down several times, tapped on the skull several times with fine forceps, and left the animal head-fixed for at least 30 mins before beginning the behavioral protocol. If necessary, we repeated this habituation protocol every day until the animal’s behavior was significant (see “Behavioral training” above). After this, we anesthetized the animal to make a small craniotomy, which was then covered with Kwik-Cast. The craniotomy was guided by fiducial marks made at the target sites for probe insertion during headplate implantation using a fine-tipped pen. Target coordinates included: AP 0.9, ML 1.7 (lAcbSh); AP 1.1 ML 1.4 (nucleus accumbens core); and AP 1.4, ML 0.6 (medial accumbens shell, mAcbSh). For the first craniotomy, a ground pin was inserted into the posterior cortex and a custom-made plastic recording chamber was fixed to the top of the headplate, both using five-minute epoxy (Devcon).

The next day, we head-fixed the mouse, covered its head as before, removed the Kwik-Cast, and flushed the craniotomy with saline. For the first recording in each craniotomy, we coated the probe in lipophilic dye at 10 mg/mL. DiI (1,1’-dioctadecyl-3,3,3’,3’-tetramethylindocarbocyanine perchlorate, Sigma-Aldrich, 42364-100MG) and DiD (1,1′-dioctadecyl-3,3,3′,3′-tetramethylindodicarbocyanine, 4-chlorobenzenesulfonate, Biotium, 60014-10mg) were dissolved in 100% ethanol (Koptec, V1001), and DiO (3,3’-dioctadecyloxacarbocyanine perchlorate, ThermoFisher, D275) was dissolved in 100% *N,N-*dimethylformamide (Sigma-Aldrich, D4254). The coated Neuropixels 1.0^147^ or four-shank Neuropixels 2.0^148^ probe was then mounted on the manipulator, and connected to the ground pin via a wire soldered onto the reference pad and shorted to ground. In the event the external reference was unstable, we used tip referencing instead. All recordings were performed in SpikeGLX software (https://github.com/billkarsh/SpikeGLX) with sampling rate = 30 kHz, LFP gain = 250, and AP gain = 500, and we analyzed only the AP channel (which was high-pass filtered in hardware with a cutoff frequency of 300 Hz).

We inserted the probe into the brain at 9 μm/s before slowing to 2 μm/s when we were 500 μm above the target depth. We stopped insertion when we saw ventral pallidal activity, characterized by large-amplitude, high-frequency spikes, on the first 40 channels or so (or 5 channels for Neuropixels 2.0). This point was usually reached around 5.2 mm below the visually-identified pial surface. After reaching the target depth, the probe was allowed to settle for 30 minutes prior to starting the experiment and Neuropixels recording. Behavioral and neural recordings were synchronized using a TTL pulse sent from the Bpod to the PXIe acquisition module SMA input at the start of every trial. After the experiment, the probe was retracted at 9 μm/s and the craniotomy was re-sealed with Kwik-Cast. Neuropixels data were spike sorted offline with Kilosort 3^149^ with default parameters, followed by manual curation in Phy (https://github.com/cortex-lab/phy).

#### Two-photon imaging

Imaging data were acquired using a custom-built two-photon microscope. A resonant scanning mirror and galvanometric mirror (Cambridge Technology, CRS 8 KHz and 6210H) separated by a scan lens-based relay on the scan head (Thorlabs, MM201) allowed fast scanning through a dichroic beamsplitter (757 nm long-pass, Semrock) and 20x/0.5 NA air immersion objective lens (Nikon, Plan Fluor). Green and red emission light were separated by a dichroic beamsplitter (568 nm long-pass, Semrock) and bandpass filters (525/50 and 641/75 nm, Semrock) and collected by GaAsP photomultiplier tubes (Hamamatsu, H7422PA-40) coupled to transimpedance amplifiers (Thorlabs, TIA60). A diode-pumped, mode-locked Ti:sapphire laser (Spectra-Physics) delivered excitation light at 920 nm with an average power of ∼60 mW at the top face of the GRIN lens^150^, modulated by a Pockels cell (Conoptics, 350-80). The microscope was controlled by ScanImage (Version 4; Vidrio Technologies). The behavior platform was mounted on an XYZ translation stage (Thorlabs, LTS150 and MLJ050) to position the mouse under the objective, and the top face of the GRIN lens was first located using a 470 nm LED (Thorlabs, M470L2).

Due to the limited axial resolution of the implanted GRIN lens, we acquired only a single imaging plane at 15.2 Hz unidirectionally with 1.4x digital zoom and a resolution of 512 x 512 pixels (∼1 μm/pixel isotropic). Imaging was either continuous or triggered 2.6 s before odor/unexpected reward onset, depending on the session. Bleaching of GCaMP7s was negligible over this time. TTL pulses were sent from the microscope to Bpod to synchronize imaging and behavioral data. Imaging typically began ∼4 weeks after GRIN implantation, to allow sufficient time for the virus to express and for inflammation to clear.

#### Two-photon pre-processing

We used the Suite2p toolbox^151^ (version 0.10.3) to register frames, detect cells, extract Ca^2+^ signals, and deconvolve these traces. We used parameter values of tau=2.0 (to approximately match the decay constant of GCaMP7f^107^), sparse_mode=False, diameter=20, high_pass=75, neucoeff=0.58; fs was set to the measured frame rate for that session (∼15.2 Hz), and all other parameters were set to their defaults. Briefly, non-rigid motion correction was used in blocks of 128 x 128 pixels to register all frames to a common reference image using phase correlation. Cell detection consisted of finding and smoothing spatial PCs and then extending ROIs spatially around the peaks in these PCs. Next, Ca^2+^ traces were extracted from each ROI after discarding any pixels belonging to multiple ROIs. Finally, neuropil contamination and deconvolved spikes were estimated in a single step from Ca^2+^ fluorescence in each ROI using the OASIS algorithm^152^ with a non-negativity constraint. This deconvolved activity was used for all subsequent analysis. ROIs were manually curated on the basis of anatomical and functional criteria using the Suite2p GUI to exclude neuropil and ROIs with few or ill-formed transients.

#### Face and body imaging

In addition to the lick port, we monitored behavior using two cameras at 30 Hz, one pointed at the face (PointGrey, FL3-U3-13Y3M) and one pointed at the body (PointGrey, CM3-U3-13S2C) under both visible and infrared LED illumination. Cameras were synchronized from Bpod once per trial using GPIO inputs, and data were written to disk via Bonsai^153^. Behavioral features were extracted using custom code alongside Facemap^92^ (version 0.2.0). Face motion energy was computed as the absolute value of the difference between consecutive frames and summed across all pixels to yield the “whisking” signal. In addition, we performed singular value decomposition (SVD) on the motion energy video (in chunks, following ref.^92^) and projected the movie onto the top 50 components to obtain their activity patterns over time. Pupil area was estimated simply as the mean (inverse) pixel value within a mask, after interpolating over blink events. Running was computed using the phase correlation of the cropped body video, to take into account limb and tail movements.

#### Optogenetic manipulation

473 nm laser light (Laserglow Technologies, LRS-0473-GFM-00100-03) was delivered to the implanted tapered fibers using a custom-built rig (modeled after refs.^154,155^) coupled to a high-performance patch cord (0.66 NA, Plexon, OPT/PC-FC-LCF-200/230-HP-2.2L KIT). Briefly, light was split into two identical paths using a 50/50 beamsplitter cube (Thorlabs, CCM1-BS013). Each path was then focused onto a galvanometric mirror (Novanta 6210K) and re-collimated using an achromatic doublet (Thorlabs, AC508-100-A-ML), before being focused onto the back of the patch cord using an aspheric condenser lens (Thorlabs, ACL50832U). This setup allowed us to modulate the angle at which light entered the patch cord, and thus the distance at which it exited the tapered fiber. We delivered light at two different angles (three in some experiments), but here we analyze only ventral manipulation trials, in which the incident angle of light was ∼0°, light exited near the tip of the fiber, and coupling between the patch cord and fiber was approximately 50%^154^.

The laser output (and the angle of the galvanometric mirrors) was controlled by Bpod via PulsePal^156^ (Version 2; Sanworks, 1102). Stimulation was delivered bilaterally during the two second-long trace period, immediately prior to reward. For CoChR excitation experiments, we used 10 ms pulses at 20 Hz with an output power at the tapered fiber of 100 μW. For GtACR1 inhibition, we used a constant, 1 mW pulse for the full 2 seconds. In both cases, stimulation was delivered on 45.5% of trials, uniformly at random across manipulation locations and trial types.

#### Histology and immunohistochemistry

Mice were deeply anesthetized with ketamine/dexmedetomidine (80/1.1 mg/kg) and then transcardially perfused using 4% paraformaldehyde. The brains were sliced at 100 μm into coronal sections using a vibratome (Leica) and stored in PBS. If performing immunostaining, slice thickness was 75 μm. These slices were then permeabilized with 0.5% triton X-100, blocked with 10% FBS, and stained with rabbit anti-tyrosine hydroxylase antibody (TH; AB152, EMD Millipore, RRID: AB_390204) at 1:750 dilution at 4°C for 24 hours to reveal dopamine axons in the striatum. Next, slices were stained with fluorescent secondary antibodies (Alexa Fluor 488 goat anti-rabbit secondary antibody, A-11008, Invitrogen, RRID: AB_143165) and DAPI at 1:500 dilution at 4°C for 24 hours. Slices were then mounted on glass slides (VECTASHIELD antifade mounting medium, H-1000, or with DAPI for non-stained slices, H-1200, Vector Laboratories) and imaged using Zeiss Axio Scan Z1 slide scanner fluorescence microscope. We visually verified the placement of all GRIN lenses and fibers to be within the lAcbSh.

### Data Analysis

#### Atlas registration

For electrophysiology experiments, we registered slices to the Allen Mouse Brain Atlas with SHARP-Track^157^ and used it to trace dyed probe trajectories in the AP and ML directions as well as visualize the registered trajectories as a coronal stack. We also used this registration to define the unique DV extent of each mouse’s lateral ventral striatal 6-OHDA lesion, and we considered only neurons that fell within this range to have been lesioned. To more accurately ascertain the depth of recordings, we used the International Brain Lab’s Ephys Atlas GUI (https://github.com/int-brain-lab/iblapps/tree/master/atlaselectrophysiology), focusing on the boundary between the ventral pallidum and nucleus accumbens due to the abrupt change in electrophysiological characteristics at this interface. When necessary, we also adopted their convention that in Allen Common Coordinate Framework^158^ (CCF) coordinates, bregma = 5400 AP, 332 DV, and 5739 ML. For plotting probe trajectories in 3D, we used the Brainrender library^159^.

For more fine-grained analysis of subregions, we used the Kim Lab atlas^160^ accessed through the BrainGlobe Atlas API^161^. This atlas applies the Franklin and Paxinos^162^ labels to the Allen CCF^158^, with additional striatal subregions defined by Hintiryan et al.^163^. For some subregions, the parcellation was finer than we needed, so we pooled subregions as follows:

● Olfactory tubercle (OT): Tu1; Tu2; Tu3
● Ventral pallidum (VP): VP
● Medial nucleus accumbens shell (mAcbSh): AcbSh
● Lateral nucleus accumbens shell (lAcbSh): lAcbSh; CB; IPACL
● Nucleus accumbens core (core): AcbC
● Ventromedial striatum (VMS): CPr, imv; CPi, vm, vm; CPi, vm, v; CPi, vm, cvm
● Ventrolateral striatum (VLS): CPr, l, vm; CPi, vl, imv; CPi, vl, v; CPi, vl, vt; CPi, vl, cvl
● Dorsomedial striatum (DMS): CPr, m; CPr, imd; CPi, dm, dl; CPi, dm, im; CPi, dm, cd; CPi, dm, dt
● Dorsolateral striatum (DLS): CPr, l, ls; CPi, dl, d; CPi, dl, imd

#### Unit inclusion criteria

To be included for analysis, units from Neuropixels recordings had to have a minimum firing rate of 0.1 Hz and to have been stable, defined as a coefficient of variation of firing rate (computed in 10 equally-sized, contiguous, disjoint blocks during the session) less than 1. 13,997 single units survived these inclusion criteria in the main dataset. In the lesion dataset, we additionally filtered neurons by their DV position: only those that fell within the DV range of the lesion were included in the matched control dataset for that mouse. Of the 9,081 neurons that survived the electrophysiological criteria, 4,879 were in the correct anatomical location, of which 2,283 came from the control and 2,596 came from the lesioned hemisphere.

#### Putative cell type identification

We assigned units to putative cell types using previously-established criteria^164^. Briefly, to be considered MSNs, units were required to have broad waveforms (Kilosort template trough-to-peak waveform duration > 400 μs) and post-spike suppression ≤ 40 ms. For the latter, we used the autocorrelation function with a bin width of 1 ms. Post-spike suppression was quantified as the duration for which the autocorrelation function was less than its average during lags between 600-900 ms.

#### Statistical software

All statistical analysis, except where explicitly stated, was performed in Python using the NumPy (v. 1.22.3), SciPy (v. 1.7.3), pandas (v. 1.1.4), scikit-learn (v. 1.0.2), statsmodels (v. 0.14.0), Matplotlib (v. 3.5.1), and seaborn (v. 0.12.2) packages^165-171^. If not otherwise specified, statistical tests used Linear Mixed Effects models (LMEs) with a random intercept for each mouse, and, if applicable, a random slope for each mouse as a function of grouping (e.g. Across-vs. Within-distribution), implemented in statsmodels. All reported *p*-values are two-tailed.

#### Units of analysis

For the behavior, control and manipulation datasets (Figs. 1, 2, 3, and 6), each observation was an individual session — that is, we used simultaneously-recorded neurons and behavior and computed effects (PCA, RDA, parallelism score, classification) on a session-by-session basis. However, given the limited spatial extent of our lesion and our lower number of simultaneously-recorded neurons, for the lesion dataset (Fig. 4) we used pseudo-populations. More specifically, we created pseudo-populations by splitting the dataset into disjoint sets of *trials*^172^, which were stitched across sessions, but not across animals. Within each session, we used simultaneously-recorded trials across neurons to preserve noise correlations where possible. For these LMEs then, pseudo-populations provided the observations and mouse was again the grouping variable. The same procedure was used for all subregion-specific analyses (Extended Data Figs. 2d, 3e, 6a-d) and ANN-based decoding (Extended Data Fig. 7a-d) due to the lower number of simultaneously-recorded neurons available for these analyses.

For the imaging dataset (Fig. 5) and ANN-based transfer (Extended Data Fig. 7e-f), we did not have enough neurons in all animals to assess distributional coding. We therefore pooled neurons not only across sessions but also across animals within genotype. Pseudo-populations were otherwise constructed exactly as in the lesion case. To be consistent with the parametric nature of LMEs while recognizing that observations were no longer specific to individual mice, we used one sample *t*-tests to assess statistical significance relative to chance levels and LMEs (with just one observation per group) to assess differences between groupings.

The only exception to these choices was when computing the fraction of cells significantly encoding each variable of interest (mean, reward, RPE, etc.), or their conjunction. In this case, we always pooled across-sessions within-mouse, since we were computing a single fraction, and used paired samples *t*-tests between data and shuffled fractions (or actual combined cells versus a prediction assuming independence).

#### Time periods for analysis

In general, we analyzed behavioral and neural data during the Late Trace period, 1-0 s before reward delivery. However, for odor decoding, we used the Odor period (0-1 s after odor onset), and reward or RPE we used the Outcome period (0-1 s after reward delivery). Neural and behavioral data were averaged within these 1 s periods before analysis, with the exception of plots of classification or regression time courses, in which averages within non-overlapping 250 ms bins were used.

#### Visualization of neural time courses

For smoothed plots of neural time courses (Figs. 1f, g; 2a; 5d, g; Extended Data Fig. 2b, 10a-b), we smoothed neural activity (spike trains or deconvolved activity traces) with a Gaussian kernel (s.d. 100 ms) before plotting or reducing dimensionality. Z-scored firing rates were computed using the mean and standard deviation of this smoothed trace. PCA time courses (Fig. 1g) were extracted by computing the average normalized, smoothed firing rate for each trial type and concatenating these into a 2D matrix of shape *N*✕(*T*✕6), where *N* is the number of neurons, *T* is the number of time points per trial, and 6 corresponds to the six possible odors. PCA was then performed and the time courses were reconstructed separately for each of the six odors. All other analyses used unsmoothed data so as to not be contaminated by later time points.

#### Principal component analysis and representational dissimilarity analysis

For two-dimensional PC plots, normalized activity during the Late Trace period was averaged across trials within a given type to produce a matrix of shape *N*✕6. We then applied PCA to reduce this matrix to shape 2✕6, having retained only the top 2 PCs. Results were qualitatively identical when using all neurons or only putative MSNs for the main dataset (Fig. 2). We report Euclidean distances between projected trial types, measured separately along each PC. RDA was similar, except that we computed cosine distances in the native (pseudo-)population normalized firing rate space, rather than a lower-dimensional projection.

#### Parallelism score

Following ref.^96^, we computed the normalized mean firing rate in response to each of the Fixed and Variable odors. There are two possible ways to pair up these four odors: (1) Fixed 1 vs. Variable 1 and Fixed 2 vs. Variable 2, or (2) Fixed 1 vs. Variable 2 and Fixed 2 vs. Variable 1. In both cases, we can compute difference vectors pointing from Variable to Fixed (Fig. 2g) and then take the cosine similarity between them. The parallelism score we report is simply this cosine similarity, averaged over the two possible divisions. Note that in the case of isotropic noise, the vectors that we define are equivalent to those defined by a maximum-margin linear classifier between the two conditions. However, high parallelism score does not necessarily imply high cross-condition generalization performance (CCGP) — for example, if the test conditions are much closer together than the training conditions, the noise is high and/or anisotropic, or the coding directions for different variables are not orthogonal (e.g. arranged as a parallelogram rather than a rectangle).

#### Classification

For both behavioral and neural binary classification, we used a support vector classifier (SVC) with a linear kernel, hinge loss function, L2 penalty, balanced accuracy scoring across classes, and regularization parameter 5 x 10^-3^, implemented in scikit-learn. The linear kernel allows for easy interpretation of the learned weights. Input data (unnormalized spike counts, lick counts, or mean Facemap predictors) were transformed using StandardScaler (computed on training data) before being fed to the classifier.

We ran five different classification analyses: CCGP^96^, pairwise decoding, congruency, mean, and odor, as described in the Main Text and figure legends. Across-distribution and within-distribution results were just the average over the relevant dichotomies (e.g. the four possible ways to set up CCGP). For all simultaneous decoding analyses except for CCGP, five cross-validation folds were used, and reported classification accuracy was the average over these five folds. For CCGP, cross-validation was unnecessary because training and test sets were fully disjoint already. Similarly, for pseudo-population based decoding (Figs. 4-5), 5 training sets and 1 disjoint test set were used in all cases. For six-way odor classification, we used multinomial logistic regression rather than SVC, again with a regularization parameter of 5 x 10^-3^ and balanced accuracy scoring across classes.

Cross-temporal decoding (Extended Data Fig. 3d, 5h-j) settings were identical to the above. For the odor, pairwise, and congruency analyses, we ensured that the same trial never appeared in both the training and testing sets, despite the different time windows used, to avoid leakage due to temporal autocorrelation. For CCGP, train and test trials were always different, so this was not a concern.

#### Cosine similarity to classification boundary

Both linear classification and regression find a high-dimensional weight vector in neural state space; computing the cosine similarity between these vectors can identify whether two analyses are honing in on the same or different features. For each session, in addition to performing classification as described above, we regressed input data (unnormalized spike counts, lick counts, or mean Facemap predictors) during the same time period against per-trial mean or variance (using StandardScaler followed by RidgeCV with default scikit-learn parameters). Note that the regression uses all six trial types, while the classification is limited to looking at only two (pairwise or CCGP) or four (congruency or mean) odors at a time. We then took the weights learned by each regression and computed the cosine similarity with the classification weights (separately for each of the five classification cross-validation folds for non-CCGP decoders; each session was summarized as the average of these five measurements). We report the results of an LME testing either the difference from a chance value of 0, indicating orthogonality (CCGP), or the difference between the absolute cosine similarities for across-and within-distribution decoders (pairwise and congruency; Extended Data Fig. 5f-g).

#### Distribution-coding subpopulation

To identify neurons that contributed significantly to distribution decoding, we extracted the coefficients from each session’s CCGP, pairwise, and congruency decoders and averaged them across dichotomies (and across cross-validation folds if necessary). For the pairwise and congruency analyses, we additionally took the difference between Across-and Within-distribution coefficients. For each quantile level (computed on each set of coefficients individually for each mouse and each decoder), we then calculated the fraction of neurons above this quantile level for all three decoders compared to null decoders in which trial types had been shuffled before being run through the decoder. We chose a cutoff such that only 2.5% of these cells from the null decoders survived; for the actual data, this corresponded to 1,600 significant distribution-coding neurons, or 11.43% of the total. We refer to these neurons as the “distribution-coding subpopulation” (Extended Data Fig. 6e-f, 7).

#### Percentage of significant cells

To compute correlations with different variables of interest, we calculated the trial-wise Pearson correlation between unsmoothed activity in a given bin and the value of the variable of interest on that trial. We then did the same thing, except that for each neuron independently we shuffled the mappings between odor and distribution. For example, when considering correlations with mean value, a Fixed 1 trial would correspond to a mean of 4 (μL). If upon shuffling, Fixed 1 odors were mapped to Nothing 2, then the corresponding mean in the shuffled dataset would be Percentages of cells significantly correlating with variables of interest (positively, negatively, or without restriction) were averaged over the four 250 ms bins corresponding to the Late Trace period, and then we subtracted the shuffled from the unshuffled fraction to account for odor coding.

#### Changes relative to Baseline

In order to assess changes in neural activity relative to the Baseline period, we first grouped all Unrewarded (Nothing) and Rewarded (Fixed and Variable) trials for each neuron. We then ran a rank-sum test between Late Trace activity and Baseline activity, separately on each neuron and trial type grouping. Finally, we computed the fraction of cells per mouse that increased or decreased significantly (α = 0.05) and then ran paired *t*-tests on the respective fractions for Rewarded versus Unrewarded trials types.

#### Comparisons across subregions, hemispheres, and genotypes

Whenever subregions, hemispheres, or genotypes were directly compared, we randomly subsampled the number of neurons so that population sizes were identical across this comparison. For subregion and hemisphere (lesioned vs. control), this matching was done within-animal. When comparing subregions, we excluded a subregion from an animal if it did not contain at least 40 neurons, hence the differing number of dots (animals) per subregion (Extended Data Fig. 2e, 3e, 6a-d, 6f. For genotype (D1 vs. D2 MSNs), matching was done across-animals, for the entire population of D1 or D2 neurons. To allow for higher neuron counts, all of these decoding analyses were performed on pseudo-populations.

#### Artificial neural network-based distribution decoding

To determine whether neural populations contained sufficient information to reconstruct the complete reward distribution, rather than simply perform binary classification based on reward variance, we constructed an artificial neural network (ANN)-based distribution decoder. Pseudo-population activity from the distribution-coding subpopulation *r* was first mapped into 16 dimensions by a trainable, unregularized decoding matrix *W*. The network takes *Wr* as input and outputs the predicted distribution. It has one input layer, two hidden layers, and one output layer. Each of the two hidden layers had 32 neurons and used the non-linear activation function *f*(*x*) =*ln*(*1* + *exp*(*x* + *1*)) − *1*, which is close to the identity function for *x* >> 0 and to-1 for *x* << 0. The output layer had size 4, with each dimension corresponding to a possible reward size (0, 2, 4, or 6 μL). After linear combination, we also applied the nonlinear function *f*(*x*) as specified above, followed by the softmax function to turn the output into a normalized probability distribution.

We applied stochastic gradient descent (SGD) to minimize the following loss function based on the 1-Wasserstein distance (D):

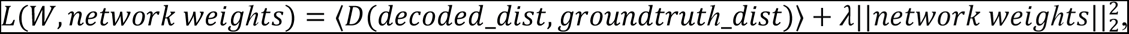

where *D* is defined as 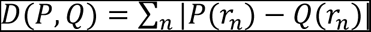 for discrete cumulative distribution functions (CDFs) *P* and *Q*, where the sum is over all used reward magnitudes, and where *r_n_* is the respective reward magnitude. In other words, the 1-Wasserstein distance measures the unsigned area between two CDFs. For plotting, we normalized this metric by dividing by the minimum achievable Wasserstein distance that would result from predicting the same distribution for every trial type across the training and test sets (“Wasserstein distance relative to reference”).

For all experiments, *λ* was set to 0.02 and the learning rate was 0.002. All the trainable weights were randomly initialized with a mean of 0 and standard deviation of 1, and then divided by 15. For each disjoint pseudo-population, we trained each of 5 candidate ANNs initialized randomly and differently for 1,200 iterations, and picked the best-performing one to further train for 10,000 iterations. The ANN was implemented in Julia (v. 1.6.7) and trained on a GPU (NVIDIA, GeForce RTX 2070).

In the standard decoding setting, all six trial types were included in the training and testing sets (with different trials in each). For decoding restricted to trial types with the same mean, only Fixed and Variable trial types were used, but split according to the same logic. In both cases, we performed decoding independently from each mouse, and we compared our results to what happened when we randomly shuffled the odor-distribution mappings before training. If merely odor identity (or, in the restricted case, mean) is encoded, then the ordered and shuffled networks should attain similar performance.

Finally, in the transfer analysis, in a similar spirit to CCGP, we trained on only four trial types and then tested on the held-out two trial types. “Matched” transfers used one Fixed and one Variable odor in the training set, assigned to the proper distribution, and evaluated performance on the corresponding test odor. “Mismatched” transfers used either two Fixed or two Variable odors in the training set, assigning one to each distribution, and evaluated performance on the held-out odors, again assigning one to each distribution. Nothing trial types were always assigned to Nothing distributions. To gain statistical power, we pooled neurons across mice for these analyses.

### Computational Modeling

In this section, we briefly review the theory behind various distributional RL algorithms before specifying the details of our implementation, for the purpose of comparing the learned code to neural activity and generating predictions for optogenetic perturbations. All models were trained for 2,000 trials per distribution.

#### Reflected expectile distributional RL (REDRL)

EDRL was first put forward as a novel machine learning algorithm^76^ and later used to explain dopamine neuron diversity in the mammalian midbrain^4^. EDRL approximately minimizes the expectile regression loss function (ER):

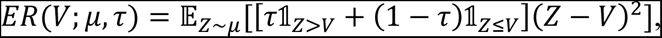

where *V* is the value predictor, *μ* is the target distribution, *Z* is a random sample from *μ*, *τ* is the asymmetry, and 1 is the indicator function, which is 1 when the subscript is satisfied and 0 when it is violated. It is an asymmetrically-weighted squared error loss function; in this sense, it generalizes the mean (squared error loss, equivalent to the 0.5th expectile) just as quantiles generalize the median^97^.

EDRL and REDRL minimize this ER loss function simultaneously for many values of *τ*, indexed by *i*, generally using SGD with respect to the value predictors (or their parameters). This formulation is sufficiently general that it can be combined with nonlinear function approximation and temporal difference learning methods, and its effectiveness has been demonstrated on the suite of Atari video games^76^. However, for simplicity, here we present the Rescorla-Wagner^173^ version of the update rule for tabular states, so the random sample from *μ* reduces to simply the reward, *r*. This is the learning rule depicted in Fig. 3:

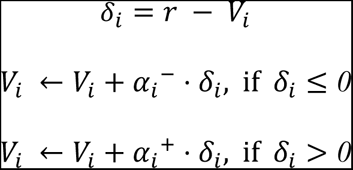

For the learning simulations (Fig. 3a), we used *α* = *α_i_^+^ + α_i_^-^* = 0.03 and initialized all value predictors to 2.

In the biological implementation of the REDRL algorithm (Fig. 3d-g), we decompose this update into two piecewise linear functions. The first function models dopamine RPEs, which are allowed to take on different slopes in the positive and negative domains, *α′_i_*^+^ and *α′_i_*^-^. The second function differs between D1 and D2 MSNs (indexed by *m*) by a reflection over the y-axis. It maps changes in dopamine firing into changes in synaptic weights^39^, which we’ll parameterize here by *β_m_* ^-/+^ (equal to 0.75/3 for D1 and 3/0.75 for D2 MSNs for the purpose of Fig. 3).

Composing these functions gives rise to the following update rules:

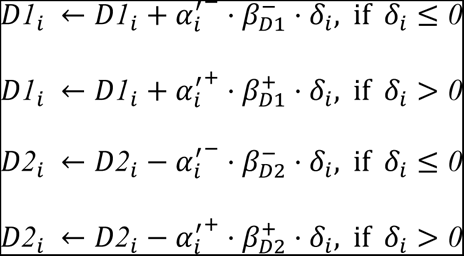

Note that D1 and D2 neurons receive unique indices *i*, so there is no overlap in the idealized case. As a consequence of the opponent plasticity rule, changes in synaptic weights in D1 and D2 MSNs have opposing effects on the encoded value predictor, modeled simply by the identity function (for D1 MSNs) or its negation, (for D2 MSNs). Therefore, this update rule becomes equivalent to the algorithmic rule if we let 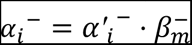 and 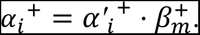 The degree of optimism or pessimism is parameterized by the dimensionless quantity 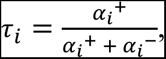 which ranges from 0 to 1. Importantly, *τ_i_* uses the net asymmetries learned by the MSNs as opposed to the asymmetries of the dopamine neurons. Therefore, both the expectile that is learned in the striatum and the zero-crossing point of the corresponding dopamine neuron are dictated by *τ_i_*, which can give rise to multiple dopamine neurons with the same apparent asymmetry but different zero-crossing points. This stands in contrast to the EDRL model, in which the dopamine neuron asymmetries alone fully determine the zero-crossing point, but nonetheless predicts the observed correlation between zero-crossing points and asymmetries^4^.

For D1 MSNs *β_m_* ^+^ > *β_m_* ^-^ and so *τ_i_* skews optimistic; analogously, for D2 MSNs *β_m_* ^+^ < *β_m_* ^-^, and *τ_i_* skews pessimistic. The precise distribution of *τ*’s will depend on the distribution of dopamine neuron asymmetries (*α′_i_*^+^ and *α′_i_*^-^) as well as the ratio of *β_m_* ^+^ to *β_m_* ^-^, neither of which has been measured precisely. To avoid making too many assumptions and to simplify interpretation, we plotted all REDRL results based on a simulation of 10 predictors with uniform spacing of *τ_i_* between 0.05 and 0.95, with all *τ_i_* > 0.5 assigned to D1 MSNs and all *τ_i_* < 0.5 assigned to D2 MSNs. Furthermore, we directly computed the expectiles of the relevant reward distributions (rather than obtaining them incrementally from samples and updates) in order to eliminate noise. We confirmed that all of our main results were robust to these choices of *τ* and simulation approach.

#### Quantile distributional RL (QDRL)

QDRL is exactly akin to EDRL, except that we minimize the quantile regression (QR) loss^72^:

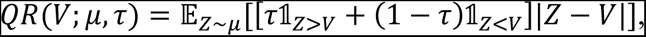

This is an asymmetrically-weighted absolute value loss function, which would return the median when positive and negative errors are balanced (*τ* = 0.5). The update rule, derived by SGD, utilizes only the sign of the prediction error, not its magnitude^97^:

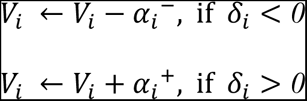

Unlike expectiles, quantiles have an intuitive interpretation: the *τ*-th quantile is the number such that *τ* fraction of samples from the distribution fall below that value and 1 - *τ* fall above it. It is therefore the inverse of the cumulative distribution function (CDF). We additionally implemented a “reflected” version of QDRL by applying the same transformation to D2 MSNs, those predictors with *τ_i_* < 0.5.

We also note that it is possible to interpolate between EDRL and QDRL using Huber quantiles^72,174^. This is simply an asymmetric squared loss within a certain interval (controlled by a hyperparameter *κ*), and a standard quantile loss outside this interval. The update rule is likewise a combination of EDRL and QDRL: piecewise linear within some range before saturating. This rule would obtain if, for example, plasticity could only change some maximum amount in either direction at any given time, as is likely the case in the brain. Notably, the Huber quantile loss is frequently used in machine learning applications^72^.

#### Categorical distributional RL (CDRL)

CDRL^71^ adopts a very different approach to learning the reward distribution. Rather than a quantile or expectile function, CDRL imagines a set of “atoms”, which function similarly to bins of a histogram. For that reason, we model these “categorical codes” using one hypothetical neuron per reward size (0-8 μL), in increments of 2 μL. The height of that bin is then assumed to be linearly (and positively) related to the firing rate of that neuron. Generalizing this scheme to use basis functions over bin values does not qualitatively alter the predictions.

#### Laplace and cumulative code

The Laplace code^83^ grew out of an effort to devise a fully local temporal difference (TD) learning rule for distributional RL. Its teaching signal is simply a sigmoidal function of reward: if reward exceeds some threshold, the neuron fires, and thresholds are heterogeneous across neurons. In the limit of infinitely steep sigmoids (Heaviside step functions), the value predictors converge to the probability that the reward exceeds the given threshold (discounted and summed over future time steps, in the TD case). This exceedance probability is equal to 1 - CDF of the reward distribution, for our simplified Rescorla-Wagner setting. By analogy to CDRL, we chose to model neural activity as linearly and positively related to this value of 1 - CDF at each of the reward bins. For completeness, we also investigated a “cumulative” code, which was just the CDF at each reward bin, or 1 - the Laplace code. The spatial derivative of this cumulative code is then equivalent to the categorical code, assuming sufficient support.

#### Actor Uncertainty (AU) model

The AU model^66^ manages to learn about reward uncertainty using biologically-plausible learning rules in D1 and D2 MSNs. We therefore wanted to test its predictions against these other models. The AU model makes use of two value predictors: one D1 and one D2 MSN, which learn as follows:

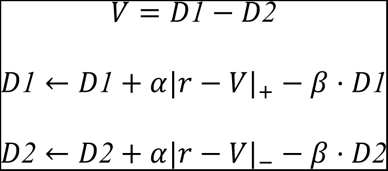

Here, |*x*|_+_ = *max*(*x*, *0*) and |*x*|_−_ = *max*(−*x*, *0*), and 0 < *β* < 1 scales the decay term to ensure stability. Using this model, it can be shown^66^ that *D1* - *D2* encodes an estimate of mean reward, and *D1* + *D2* encodes an estimate of reward spread. For our implementation, we set *α* = 0.1 and *β* = 0.01.

#### Distributed AU model

The distributed AU model^175^ works similarly, except that we now allow there to be different learning rates *α_i_^+^* and *α_i_^-^* for D1 and D2 MSNs, respectively, just as in the distributional RL setting. The difference *V_i_ = D1_i_ - D2_i_* approximates the *τ_i_*-th expectile, biased by *β*. For our simulations, we chose *α* = *α_i_^+^ + α_i_^-^* = 0.2 and *β* = 0.01.

#### Modeling perturbations

Simulating optogenetic inhibition and excitation in these models (Extended Data Fig. 11) required slightly different choices, depending on the type of code. For expectile, quantile, and AU-based models, we clamped the relevant simulated neuron(s) to either 0 or 8, the maximum reward value across all distributions, to simulate model inhibition and excitation, respectively. Note that it was the neural activity (*D1_i_* or *D2_i_*) that we were directly clamping when applicable, not the value prediction it encoded (*V_i_*). For the expectile and quantile models, optimistic and pessimistic perturbations meant clamping the value of predictors with *τ_i_* > 0.5 and *τ_i_* < 0.5 respectively. For the AU model, they were identified with the D1 and D2 MSN, respectively.

Finally, for the distributed AU model, we implemented two versions of the perturbation, one in which all D1 (optimistic) or all D2 (pessimistic) neurons were manipulated, and one in which only those with *τ_i_* > 0.5 or *τ_i_* < 0.5, respectively, were manipulated. We call the latter the “Partial Distributed AU” model, for the purposes of model comparison. For the AU models, it is only the difference *D1_i_ - D2_i_* that is bounded within the range of reward sizes, not the activities individually. We therefore added or subtracted a fixed amount (the maximum reward size across all trial types, 8 μL) across reward predictors to simulate excitation or inhibition, respectively, in these models, rather than clamping their value to a constant.

For categorical, cumulative, and Laplace codes, the semantics of each simulated neuron are different: their activations range from 0 to 1 and encode a (cumulative) probability, rather than a value. Thus, inhibiting or exciting them meant changing the relevant probability to 0 or 1, respectively. Pessimistic neurons were those that corresponded to the 0 or 2 μL bins, and optimistic neurons corresponded to 6 and 8 μL. To reconstitute a properly-normalized probability distribution after the perturbation, in the case of the categorical code, we divided by the sum of the predictors (or made it a uniform distribution if the sum was zero). For the categorical and Laplace codes, we took the spatial derivative of the implied CDF, subtracted off the minimum if any value was negative, and then divided by the sum (or made it uniform if the sum was zero).

In all cases, we found the mean of the (imputed) perturbed probability distribution and then compared it to the mean without any perturbation to model the effect of optogenetic manipulation on lick rate.

#### Model comparison

We used the predicted Manipulation - No Manipulation differences from each model as a regressor with which to predict the difference in licking across trial types, averaged across mice, using linear regression (with no intercept term). Separate regressions were fit for inhibition and excitation to allow for potentially different scaling in each case, and their coefficients of determination were averaged to produce a single summary measure of goodness of fit.

#### Data availability

Pre-processed data will be posted to online repositories upon publication.

#### Code availability

Analysis code will be posted to online repositories upon publication.

## Acknowledgments

We thank members of the Uchida Lab for valuable discussions and comments on the manuscript. Ed Soucy and Brett Graham of the Harvard Center for Brain Science Neurotechnology Core Facility provided critical assistance with instrumentation. We’d also like to thank Dr. Allison Girasole and Prof. Bernardo Sabatini for sharing the GtACR1 mouse line; Dr. Xintong Cai, Prof. Bernardo Sabatini, Prof. Chris Harvey, and Prof. Sam Gershman for helpful conversations; and Dr. Matteo Carandini, Dr. Kenneth Harris, Dr. Andrew Peters and other members of the Cortex lab for their advice on Neuropixels recording. This work was supported by grants from NIH (R01NS116753, to N.U. and J.D.; F31NS124095, to A.S.L.), the Human Frontier Science Program (LT000801/2018, to S.M.), the Harvard Brain Science Initiative, and the Brain and Behavior Research Foundation (NARSAD Young Investigator no. 30035 to S.M.). We thank the Harvard Center for Biological Imaging (RRID:SCR_018673) for infrastructure and support for *ex vivo* imaging, which was funded in part by the Simmons Award (to A.S.L.). The computations in this paper were run in part on the FASRC Cannon cluster supported by the FAS Division of Science Research Computing Group at Harvard University.

## Author Contributions

A.S.L. and N.U. designed the experiments. A.S.L. and M.M. performed the experiments, with initial help from S.M. A.S.L. and M.M. preprocessed the data. A.S.L. analyzed the data and implemented the computational models with input from J.D. and N.U. Q.Z. implemented ANN-based distributional decoding under the supervision of J.D. A.S.L. wrote the first draft of the manuscript and created the figures. N.U., J.D., S.M., and A.S.L. edited the manuscript.

## Materials and Correspondence

Please direct any requests for materials to Naoshige Uchida.

---

**Extended Data Fig. 1.**
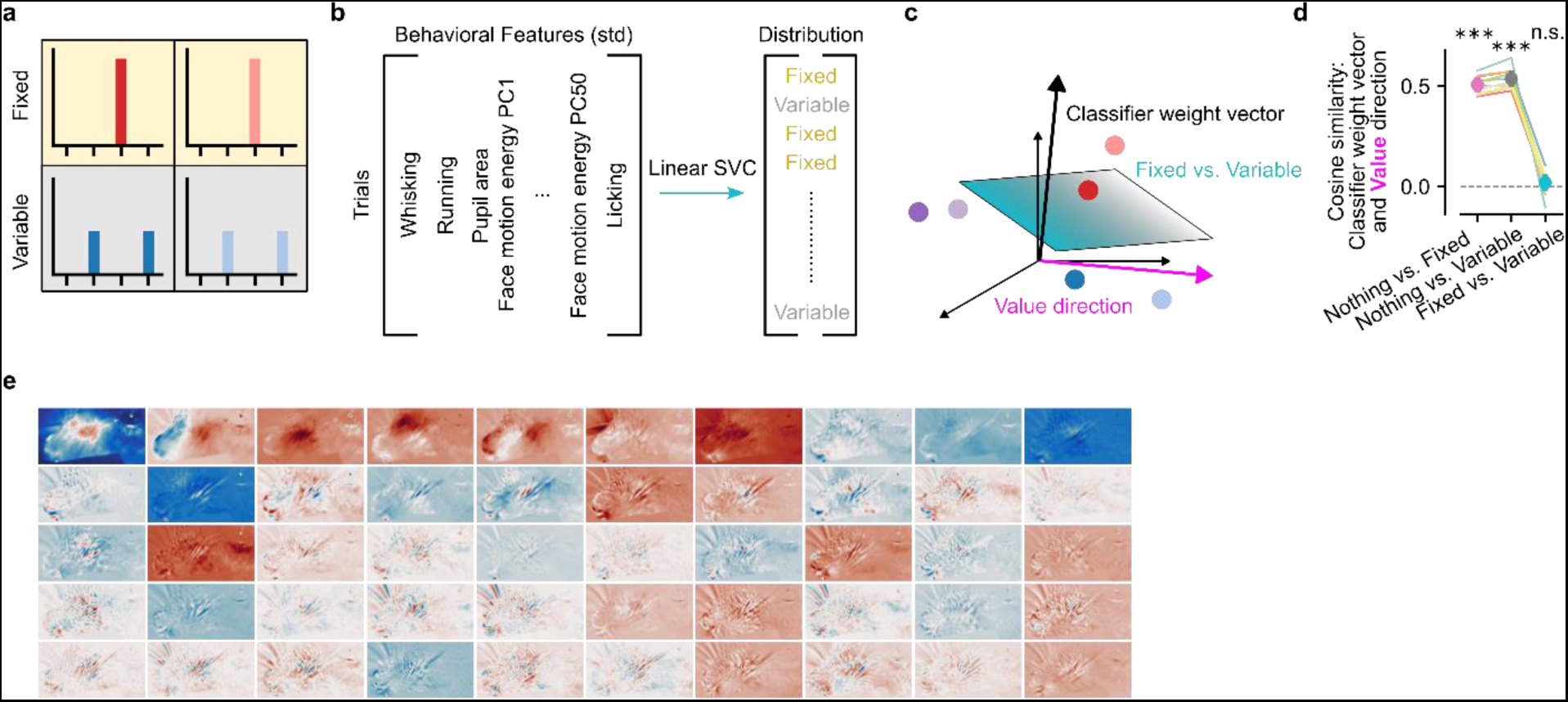
**| Behavioral classification analysis. a**, Odors corresponding to the same distribution were treated as the same class. This is illustrated for the case of Fixed vs. Variable classification, with the background shading (yellow vs. grey) indicating the target for the classifier. **b**, Schematic of behavioral classification. On each validation fold, whisking, running, pupil area, licking, and the top 50 face motion energy PCs in the training set were z-scored and then passed to a support vector classifier (SVC) with a linear kernel, which predicts the associated distribution. **c**, Schematic of orthogonality analysis. The weights learned by the SVC define a vector orthogonal to the hyperplane that best separates distributions. A separate vector can be defined by regressing the mean reward (“Value direction”) of each trial against their corresponding behavioral regressors. While the SVC hyperplane considers only four odors at a time, the regression direction takes into account all six odors. **d**, Cosine similarity between the classifier weight vector and the Value direction. Any differences in behavior between Fixed and Variable trials are orthogonal to Value (relative to chance level of 0: *p* < 0.001 for Nothing vs. Fixed, *p* < 0.001 for Nothing vs. Variable, *p* = 0.154 for Fixed vs. Variable). **e**, Spatial masks corresponding to face motion energy PCs in an example session, sorted by variance explained. Successive PCs emphasize finer and finer aspects of mouse whisking, sniffing, and licking behavior.

**Extended Data Fig. 2.**
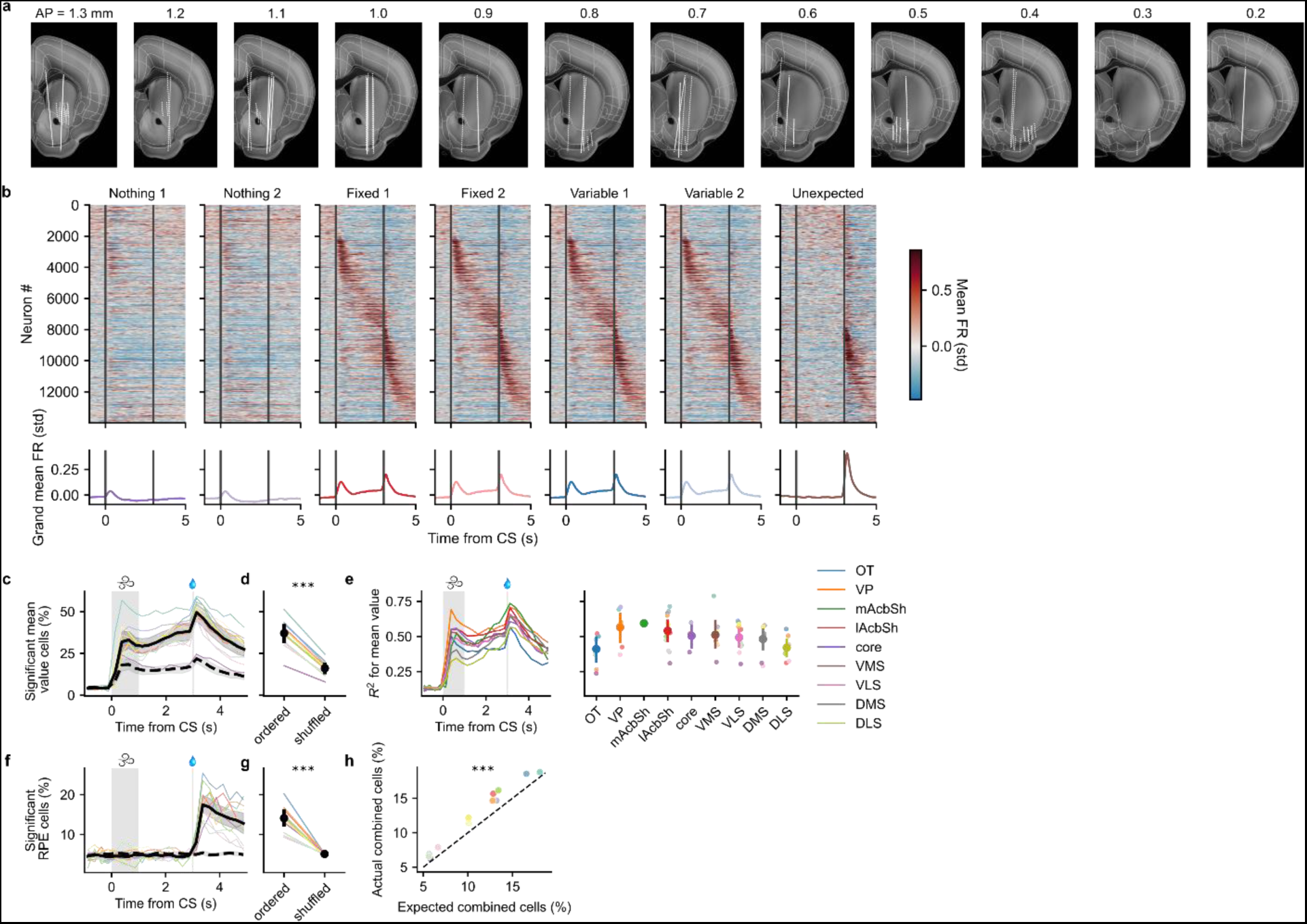
**| Value and RPE coding across the striatum**. **a**, Serial coronal sections showing recording sites of probe insertions (white dotted lines), registered to the Allen Common Coordinate Framework. **b**, *Top*, heatmaps showing average z-scored firing rate in response to each odor for each neuron. Neurons were sorted according to the time of peak activity when averaged on half of Variable 2 odor trials, and then plotted in this same order for the remainder of trials, grouped by trial type. The seventh and final trial type corresponds to Unexpected rewards, which were not preceded by an odor. *Bottom*, grand average z-scored firing rate across all neurons. **c**, Fraction of neurons that significantly correlate with mean reward, computed separately in non-overlapping 250 ms time bins. Each mouse is shown in a different color, with the mean ± 95% confidence interval across mice shown in solid black. Dashed line is the average across mice after shuffling the mapping between odors and distributions, thereby accounting for pure odor coding. **d**, Average percentage of significant cells during the Late Trace period (*p* < 0.001, paired samples *t*-test). **e**, *Left*, cross-validated *R*^2^ predicting the mean reward on each trial as a function of striatal subregion, computed separately in non-overlapping 250 ms time bins. To ensure fair comparison across subregions, we for each animal generated multiple pseudo-populations of 40 neurons each by repeatedly sampling without replacement neural subpopulation across session boundaries until there were fewer than 40 neurons remaining. Animals with fewer than 40 neurons in the given region were excluded. Lines show averages across mice for each subregion. *Right*, average *R*^2^ over the Late Trace period. Smaller dots show averages across pseudo-populations for each mouse with at least 40 neurons in that region. **f**, Same as **c**, except showing the fraction of neurons that significantly correlate with reward prediction error (RPE), defined as the difference between actual and expected reward. **g**, Same as **d**, except showing the average percentage of significant cells during the Outcome period, 0-1 s after reward delivery (*p* < 0.001). **h**, The actual fraction of cells in each mouse that significantly correlated with both mean value and RPE was compared to the product of the individual fractions for mean and RPE-coding cells (the predicted fraction assuming independence; *p* < 0.001, paired samples *t*-test).

**Extended Data Fig. 3.**
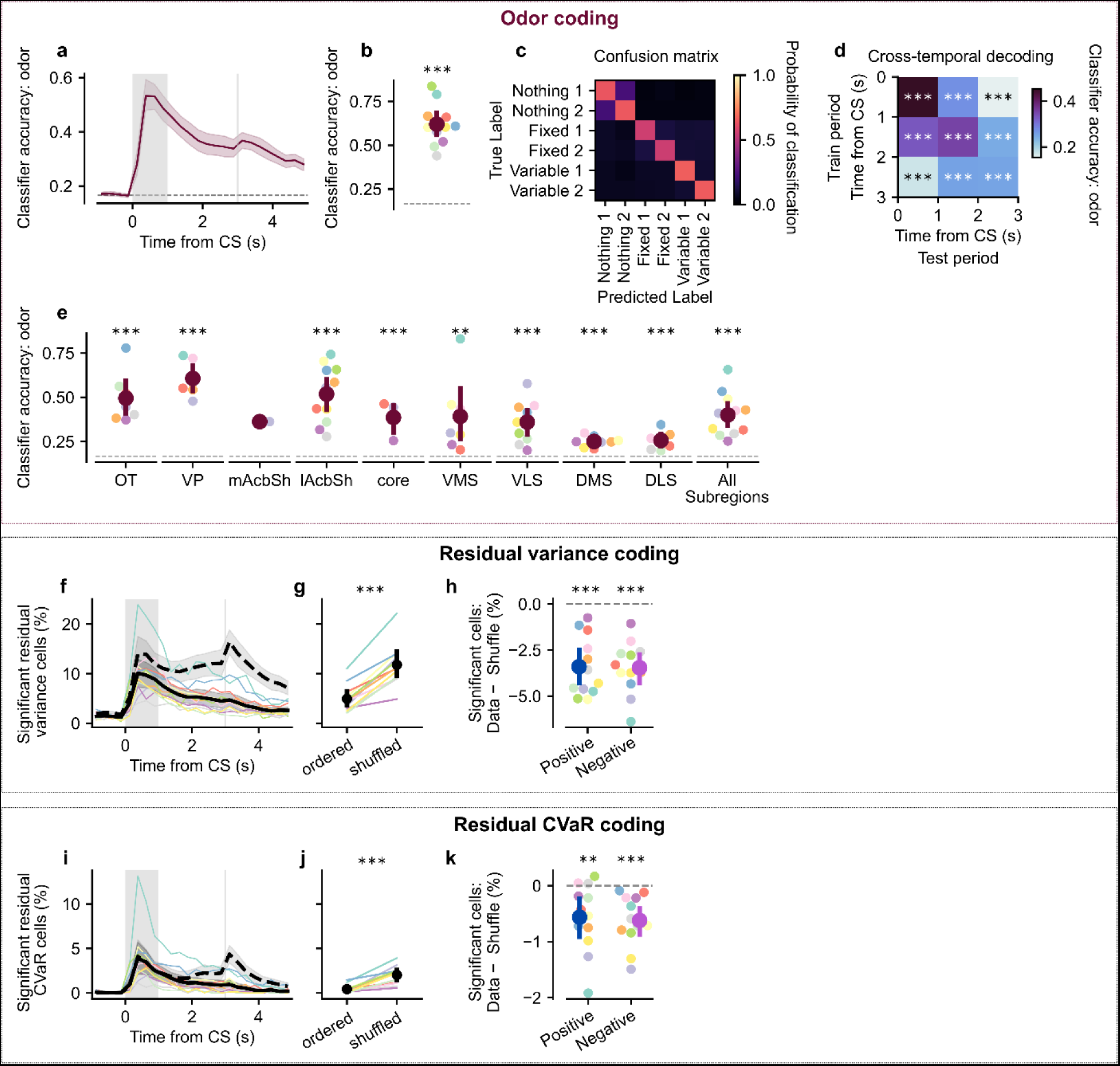
**| Odor and residual variance coding in the striatum**. **a**, Decoding accuracy across time of a multinomial logistic regression classifier decoding odor identity. **b**, Quantification of **a** during the Odor period (*p* < 0.001 relative to chance level of 1/6). **c**, Confusion matrix for odor decoding during the odor period shows high decoding accuracy for all odors, with relatively higher confusability for odors with the same mean. **d**, Cross-temporal decoding reveals that odor decoding is stable across time, allowing a classifier trained e.g. on Late Trace period activity to generalize well above chance to the Odor period, and vice versa (all *p*’s < 0.001 relative to chance level of 1/6). **e**, Pseudo-population odor decoding across subregions (see Methods section titled “Comparisons across subregions, hemispheres, and genotypes”). OT, olfactory tubercle; VP, ventral pallidum; mAcbSh, medial nucleus accumbens shell; lAcbSh, lateral nucleus accumbens shell; core, nucleus accumbens core; VMS, ventromedial striatum; VLS, ventrolateral striatum; DMS, dorsomedial striatum; DLS, dorsolateral striatum (*N* = 1 mouse for mAcbSh, *p* = 0.006 for VMS, all other *p*’s < 0.001). **f**, Same as Extended Data Fig. 2c, except showing the fraction of neurons that significantly correlate with variance, after regressing out the contribution of mean reward coding separately for each time bin. **g**, Average percentage of significant Residual Variance cells during the Late Trace period is *less* than would be predicted from odor coding alone (*p* < 0.001, paired samples *t*-test). **h**, Same as Fig. 3m, except for Residual Variance coding. Fraction is lower than chance for both positive-and negative-coding cells (*p* < 0.001, paired samples *t*-test). **i-k**, Same as **f-h**, but for conditional value at risk (CVaR), a common risk measure used in finance and reinforcement learning^126,176,177^, defined as the expected value within the lower *α*-quantile of a probability distribution. For our distributions, this will be equivalent to the mean for *α* > 0.5 and equivalent to the minimum value for *α* < 0.5, which differs only for the Variable distribution, where it is 2. The latter is what we plot here, after regressing out mean coding. Again, there are fewer Residual CVaR cells than would be expected from odor coding alone (*p* < 0.001, paired samples *t*-test) and this is true for both positive-and negative-coding cells (*p* = 0.009 and *p* < 0.001, respectively, paired samples *t*-test).

**Extended Data Fig. 4.**
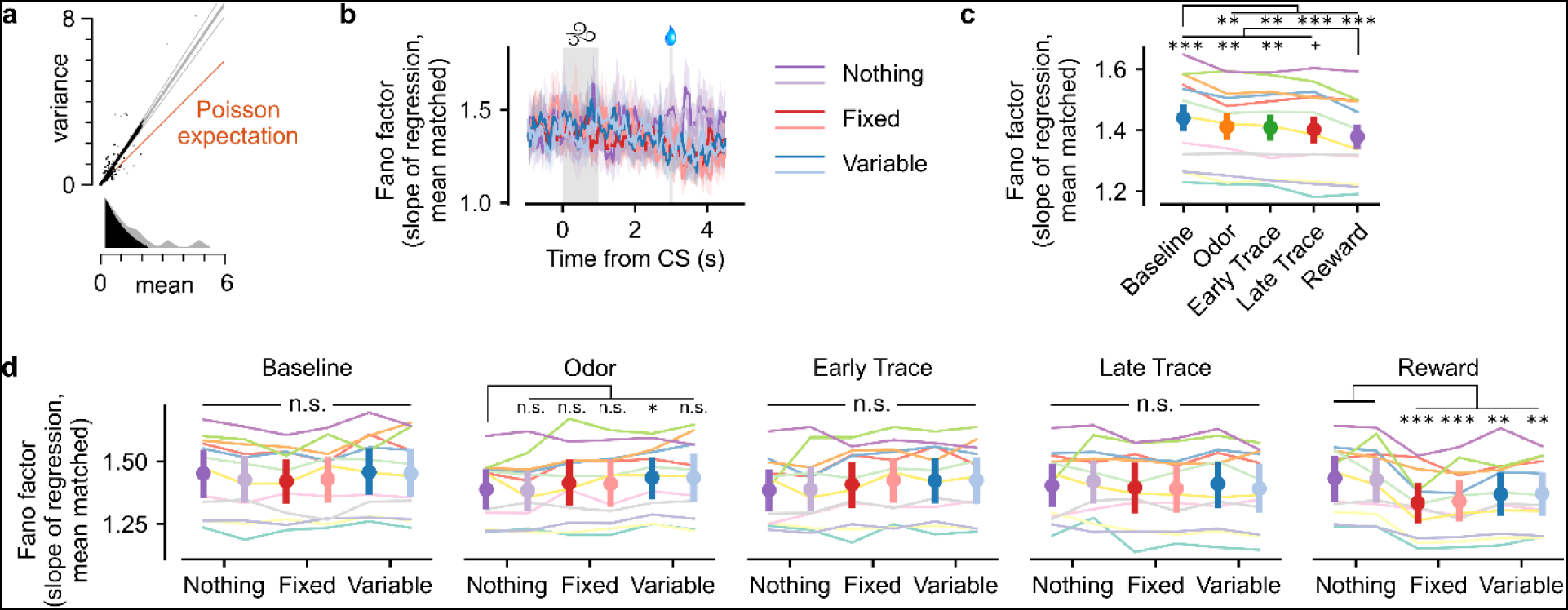
**| Sampling-based codes are inconsistent with striatal activity patterns. a**, Illustration of how the mean-matched Fano factor was computed^178^. Spike counts were computed in 100 ms bins for each trial. The mean and variance (across trials) of that count then contributed one data point to the scatter plot. Grey dots depict all neurons from an example session, time bin (centered 200 ms after odor onset), and odor (Variable 2). The grey line is the regression fit to all data, constrained to pass through zero and weighted according to the estimated s.e.m. of each variance measurement. Black dots are the data points preserved by mean matching at each time point, to eliminate the possibility that differences across time are driven by differences in firing rates, which could in principle violate the Poisson assumption. This transforms the distribution of mean counts from the grey to the black distribution. The regression slope for the mean matched data is plotted as the black line. Finally, the Poisson expectation of equal mean and variance is plotted in orange, with a slope of one. This procedure was performed independently on each session, time bin, and trial type. **b**, Time course of the computed mean-matched Fano factor (± 95% confidence interval) for the example session shown in **a**. That is, the slope of black line in **a** is the height of the light blue, Variable 2 line in **b** 200 ms after CS onset. **c**, Quantification of mean matched Fano factor across second-long time periods. Consistent with cortical observations^178^, we see a quenching of variability upon CS onset (Baseline: *p* = 0.002, 0.001, < 0.001, < 0.001 relative to Odor, Early Trace, Late Trace, and Reward periods), and another one upon reward delivery (Reward: *p* < 0.001, = 0.002, 0.006, 0.053 for Baseline, Odor, Early, and Late Trace periods). **d**, Quantification of mean matched Fano factor across trial types, shown separately for each time period. In general, there is no tendency for Variable odors to elicit strong and sustained increases in variability, as would be predicted by sampling-based codes (Baseline, Odor, Early and Late Trace: all *p*’s > 0.05, except Nothing 1 vs. Variable 1 for Odor: *p* = 0.032 uncorrected). However, reward delivery specifically drives yet another decrease in variability (Nothing 1: *p* = 0.570 for Nothing 2; *p* < 0.001 for Fixed odors; *p* = 0.002 for Variable odors).

**Extended Data Fig. 5.**
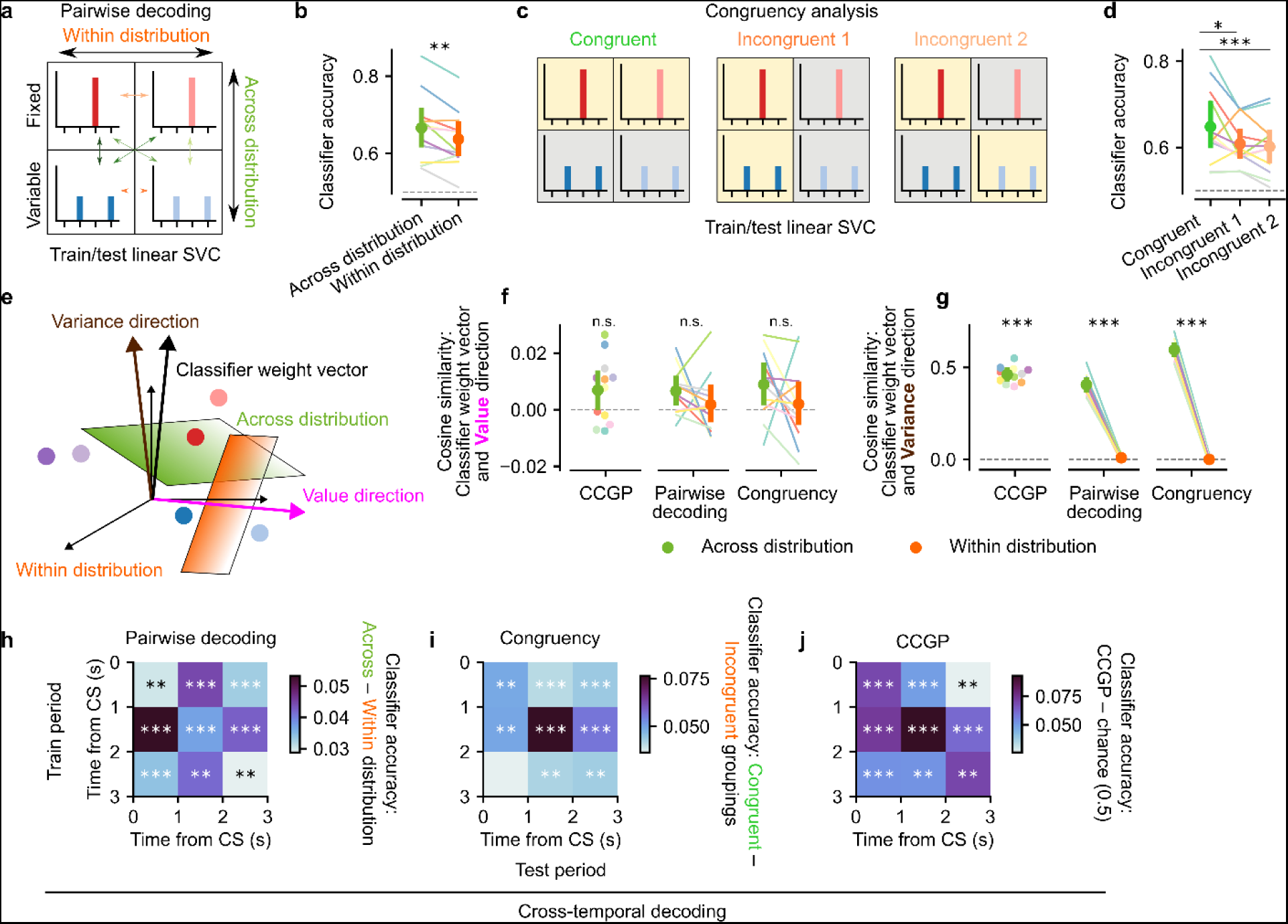
**| Distributional coding is robust, orthogonal to value, and consistent across time**. **a**, Schematic of pairwise decoding analysis. Linear SVCs were trained on individual Fixed and Variable odors, two at a time. This resulted in six possible dichotomies, four of which encompassed one Fixed and one Variable odor (green arrows; “Across distribution”) and two of which compared odors cuing the same exact distribution (orange arrows; “Within distribution”). **b**, Pairwise decoding during the Late Trace period was significantly better for across-than within-distribution pairs, consistent with distributional but not traditional RL (*p* = 0.001). **c**, Schematic of congruency analysis, which considered all four Fixed and Variable odors simultaneously. In the Congruent grouping, both Fixed odors were assigned to one class (yellow background) and both Variable odors were assigned to the other class (grey background), just as was done for behavioral decoding. By contrast, in the Incongruent groupings, class assignments cut across Fixed and Variable distributions. **d**, Classifier accuracy in the Late Trace period was higher for Congruent than Incongruent pairs, again consistent with distributional but not traditional RL (Congruent: *p* = 0.028 vs. Incongruent 1, *p* < 0.001 vs. Incongruent 2). **e**, Schematic illustrating the classifier weight vector (normal to the separating hyperplane for across-or within-distribution classifications) and the regression weight vector (for Value or Variance). **f**, Quantification of cosine similarity between the classifier weight vector and the Value direction shows that the vectors are not significantly different from orthogonal (CCGP: *p* = 0.071 relative to chance value of 0; Pairwise: *p* = 0.797 Across-vs. Within-distribution absolute cosine similarity; Congruency: *p* = 0.493 Across-vs. Within-distribution absolute cosine similarity). **g**, Same as **f**, but for Variance rather than Value direction (*p* < 0.001 for all comparisons). **h-j**, Cross-temporal decoding for the pairwise, congruency, and CCGP analyses. Distributional RL is favored during every time period between odor onset and reward delivery, and decoders trained during one period almost always generalize to other time periods.

**Extended Data Fig. 6.**
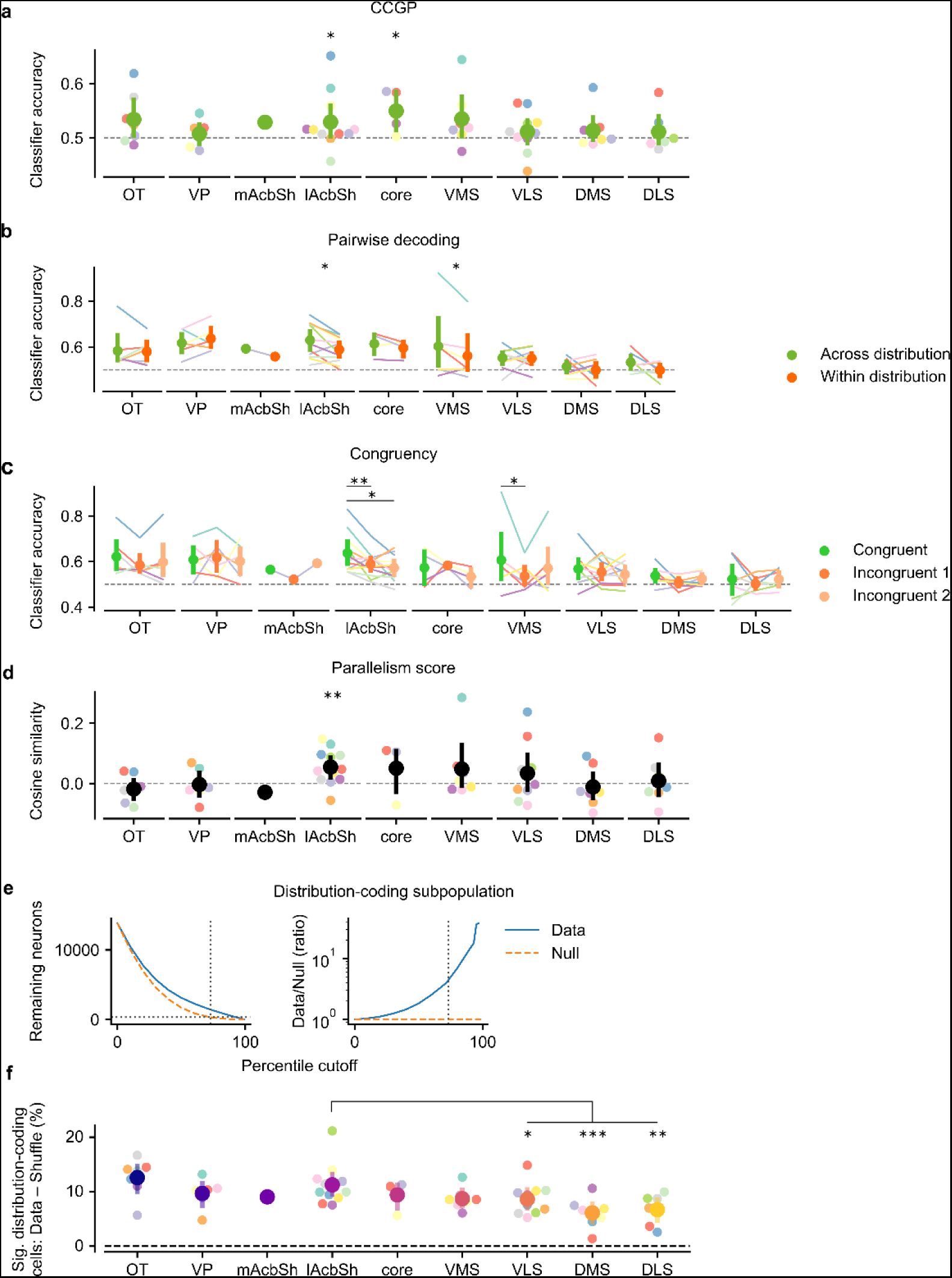
**| Distributional coding is strongest in the lAcbSh. a**, Pseudo-population CCGP across subregions (relative to chance level of 0.5: *p* = 0.059, 0.473, 0.044, 0.017, 0.088, 0.346, 0.257, 0.407, and 0.133 for OT, VP, mAcbSh, lAcbSh, core, VMS, VLS, DMS, and DLS, respectively. Same order applies to all statistics in this figure). Pseudo-populations were constructed as in Extended Data Fig. 3e. b, Pseudo-population pairwise decoding across subregions (Across-vs. Within-distribution: *p* = 0.861, 0.344, 0.883, 0.010, 0.409, 0.040, 0.882, 0.482, 0.106). **c**, Pseudo-population congruency analysis across subregions (Congruent vs. Incongruent 1: *p* = 0.097, 0.817, 0.744, 0.007, 0.832, 0.047, 0.523, 0.138, 0.523; Congruent vs. Incongruent 2: *p* = 0.306, 0.760, 0.815, 0.010, 0.473, 0.177, 0.316, 0.486, 0.985). **d**, Parallelism score across subregions (relative to chance level of 0: *p* = 0.300, 0.878, 1.00, 0.001, 0.229, 0.243, 0.273, 0.615, 0.764). **e**, *Left*, fraction of neurons with classifier coefficients above the percentile cutoff for all three (CCGP, pairwise, and congruency) analyses. Horizontal dotted line indicates level at which 2.5% of null coefficients fell above the cutoff; this was the 73rd percentile (vertical dotted line), and retained 11.43% of neurons. *Right*, ratio of data to null coefficients falling above the cutoff (log scale). **f**, Fraction of distribution-coding cells in each subregion. This fraction is significantly higher in the lAcbSh than in more dorsal subregions (relative to lAcbSh: *p* = 0.339, 0.285, 0.473, 0.274, 0.071, 0.038, 0.001 for OT, VP, mAcbSh, core, VMS, VLS, and DLS, respectively; *p* < 0.001 for DMS).

**Extended Data Fig. 7.**
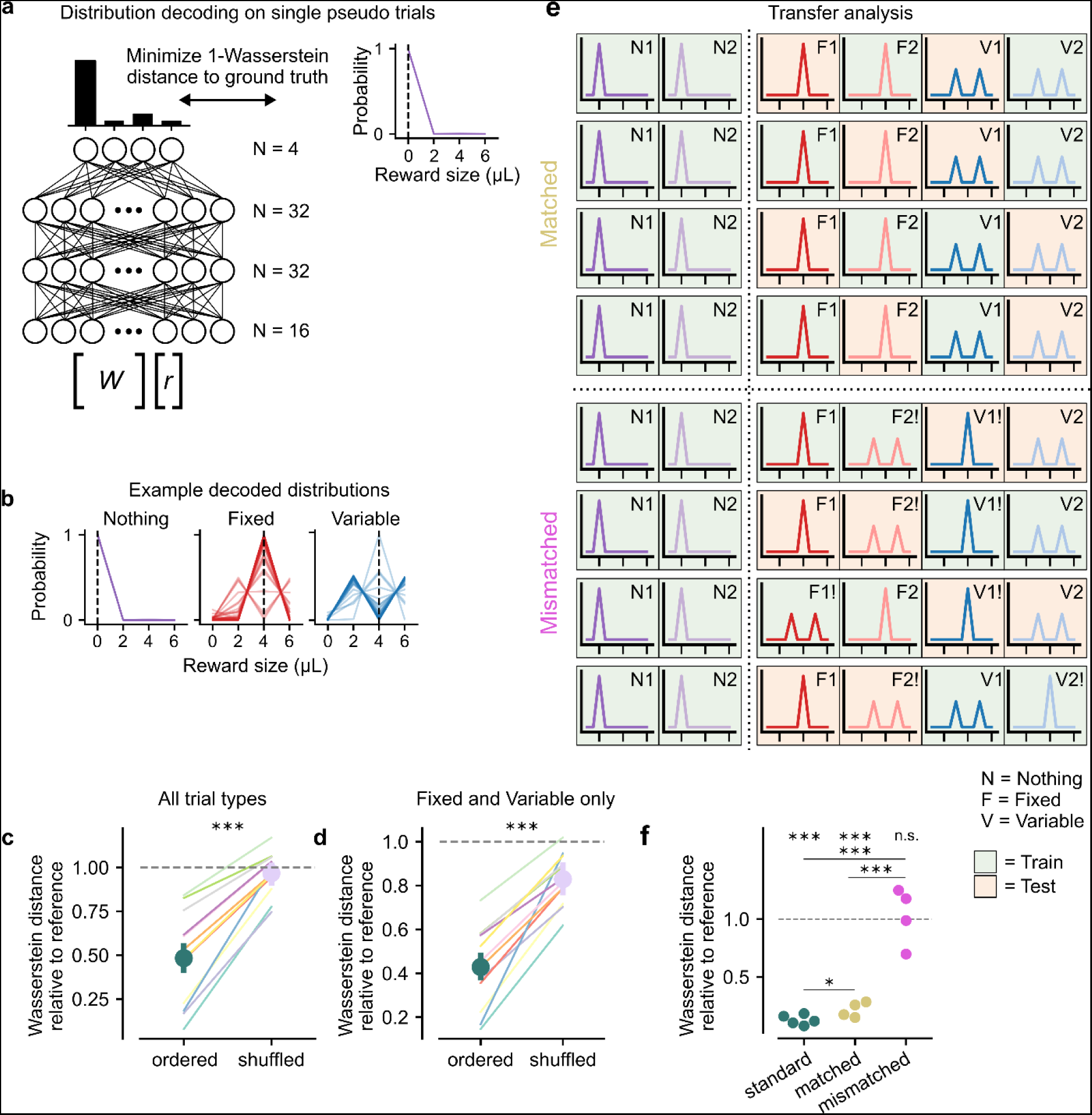
**| Artificial neural network-based distribution decoding captures information beyond the mean. a**, ANN schematic. Single-trial spike counts from the distribution-coding subpopulation *r* were linearly mapped into 16 dimensions by the trainable matrix *W* and then fed through the network (see Methods). After a final layer, a softmax function transformed activations into a properly-normalized probability distribution, whose 1-Wasserstein distance to ground truth was minimized with stochastic gradient descent. **b**, Example decoded distributions from the test set, shown as line plots to distinguish individual pseudo-trials. **c**, Wasserstein distance relative to reference for the ANN trained on all six trial types, with and without shuffling odor-distribution mappings (*p* < 0.001 ordered vs. shuffled; *p* < 0.001 ordered relative to chance value of 1; *p* = 0.350 shuffled relative to chance value of 1). **d**, Same as **c**, but for ANN trained on only the Rewarded odors, which shared the same mean (*p* < 0.001 ordered vs. shuffled, ordered relative to chance value of 1, and shuffled relative to chance value of 1). **e**, Schematic depicting setup for transfer analysis. Four trial types, including both Nothing odors, were used for training (green background), and the other two were used for testing (orange background). Matched pairings veridically assigned odors to distributions, while mismatched pairings used either only Fixed or only Variable odors for training while assigning one member per training pair and one member per testing pair to the opposite distribution (indicated by the exclamation mark). There were four possible ways to draw the matched dichotomies, all of which are shown (rows). For the mismatched dichotomies, the test labels could be flipped arbitrarily, so only one possibility (the F2 and V1 distributions swapped for testing) is shown for each training set. **f**, Wasserstein distance relative to reference for standard, matched, and mismatched settings. Standard is identical to analysis shown in **c**, except that for this decoder, neurons from all mice were pooled. Matched transfer yields distributions that are nearly as accurate as training with all six trial types (*p* < 0.001 for matched vs. mismatched and standard vs. mismatched, independent samples *t*-test; *p* = 0.043 for standard vs. matched, independent samples *t*-test; *p* < 0.001 for standard and matched relative to chance value of 1, one-sample *t*-test; *p* = 0.836 for mismatched relative to chance value of 1, one-sample *t*-test).

**Extended Data Fig. 8.**
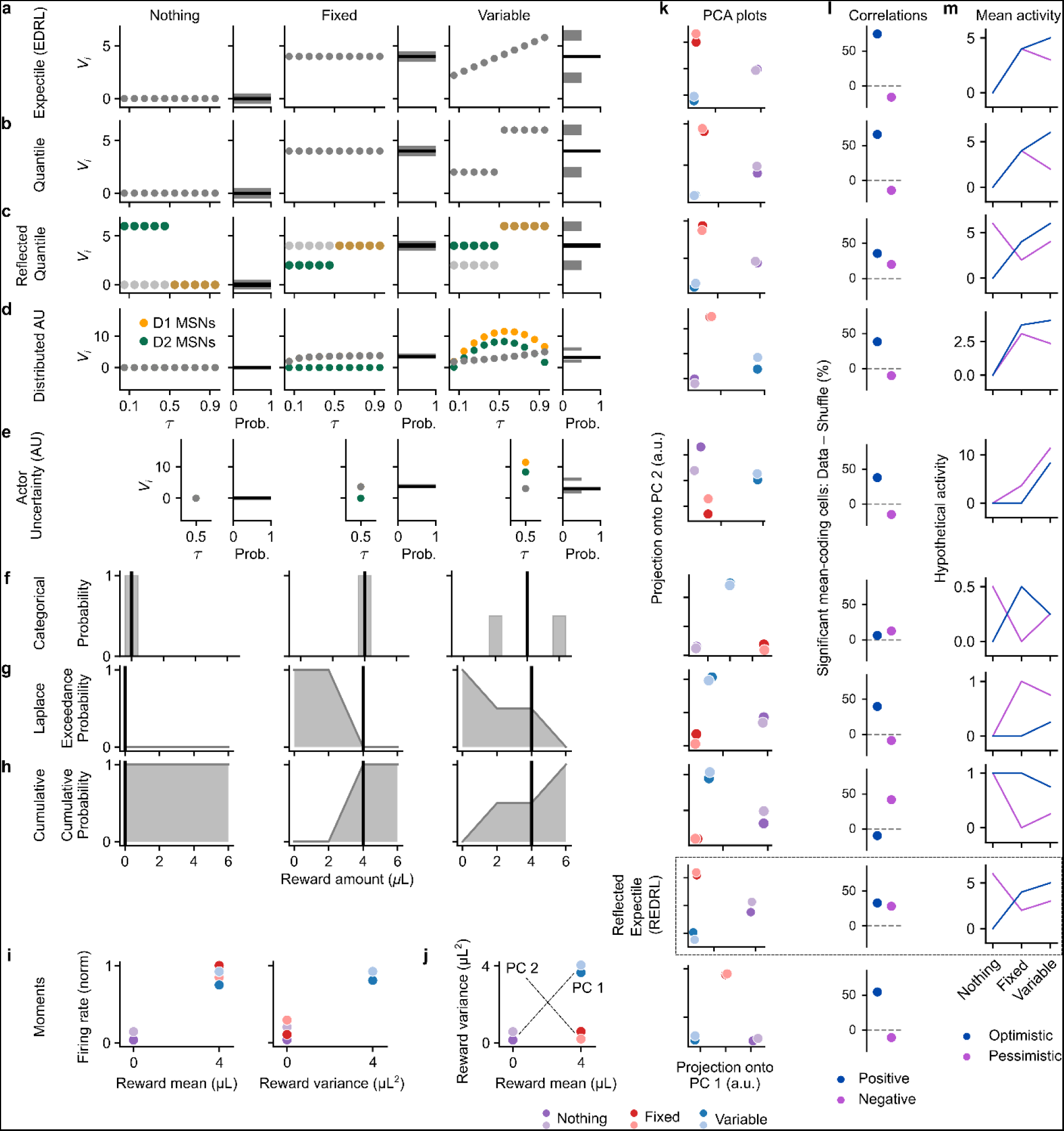
**| Additional detail for distributional model comparisons. a**, Schematic showing converged expectile code for each distribution (Nothing, Fixed, and Variable) learned by EDRL, as in Fig. 3b. The activation of each value predictor is shown as a function of *τ*, the level of pessimism or optimism. Together, they encompass the complete reward distribution. **b**, Same as **a**, but for quantiles rather than expectiles. **c**, Same as **b**, but for a reflected quantile code in which pessimistic (D2, green) neurons correlate negatively with *V_i_* (grey). Optimistic (D1, yellow) neurons are identical to *V_i_*, as in REDRL. **d**, Same as **a**, but showing the converged value predictors for the Distributed Actor Uncertainty model^175^. In it, D1 and D2 MSNs learn exclusively from positive and negative RPEs, respectively, such that their difference at each level of *τ* (grey dots) approximates each expectile, and their sum relates to the spread of the distribution. This drives maximal activity in response to Variable odors, which is why they separate out most clearly along PC 1. **e**, Same as **d**, but for a reduced version in which only a single pair of value predictors are learned with balanced positive and negative learning rates^66^ (*τ* = 0.5). **f**, Same as **a**, but for a categorical code in which distributions are encoded as a histogram^71^. Each neuron is imagined to correspond to a single reward bin, with its firing rate proportional to the height of that bin. **g**, Same as **f**, but for a Laplace code^83^. In the limit of infinitely steep reward sensitivities for the teaching signal, these value predictors converge to the probability that the reward delivered exceeds some threshold reward amount, the “exceedance probability.” This is simply 1 minus the CDF of the probability distribution in question. Neural activities are taken to be proportional to this 1 - CDF value. **h**, Same as **g**, but for a population of neurons that flips the encoding, and so is directly proportional to the CDF. **i**, A hypothetical “distributional” code in which each neuron’s firing rate linearly correlates with either reward mean (*left*) or variance (*right*). **j**, Each trial type, replotted in mean-variance space. From this picture, it is clear that for this particular set of reward distributions, Fixed odors will be located at the midpoint between Nothing and Variable odors along PC 1, though altering the ratio of mean-to variance-coding neurons will move Fixed odors left or right along PC 1. Different sets of reward distributions could lead to different geometries. **k-m**, Qualitative features of each code in **a-i** plus random noise. REDRL predictions from Fig. 3 are included in the box on the second-to-last line, for comparison. **k**, PCA projection for each code. Only quantile-like codes give rise to the pattern observed in the data. **l**, Percentage of simulated predictors that significantly correlate with mean reward either positively (blue) or negatively (purple) for each code type. Only the reflected and categorical codes have a substantial fraction of both types of cells. In practice the positive-coding predictors are optimistic and the negative-coding predictors are pessimistic. **m**, Hypothetical activity in response to each distribution, averaged separately over optimistic (blue) and pessimistic (purple) predictors for each code type. Only the reflected codes and AU model predict a noticeable uptick in Variable relative to Fixed odors.

**Extended Data Fig. 9.**
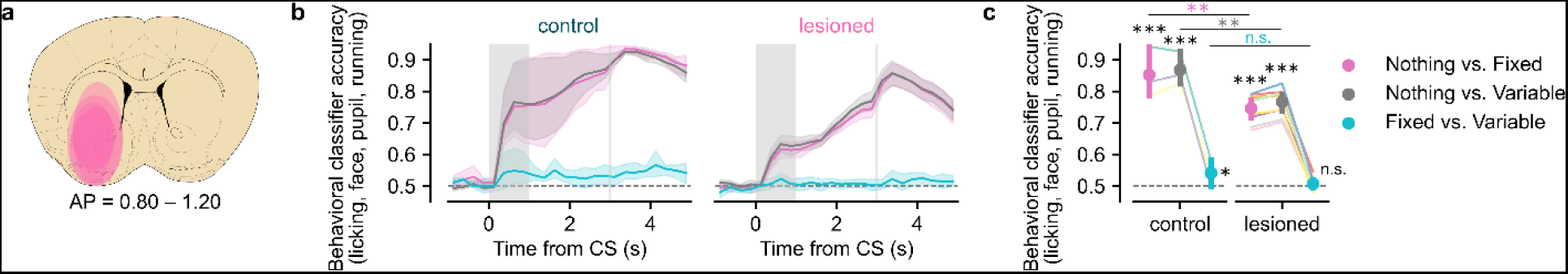
**| Quantification of 6-OHDA lesion extent, location, and behavior. a**, Consensus heat map of all five animals’ lesion locations. 6-OHDA was injected in the lAcbSh but diffused into the VLS, so we considered both regions to be lesioned. We excluded OT, despite the fact that it was often lesioned, because it is not physically contiguous and showed weaker evidence of distributional coding in control animals. **b**, Behavioral decoding analysis comparing fully intact animals (*N* = 3) and unilaterally lesioned (*N* = 9) animals across time. For this analysis, animals were considered lesioned if they had received any 6-OHDA injection, even if that hemisphere was never recorded or was mistargeted relative to Neuropixels recording location. **c**, Quantification of behavioral classifier accuracy during the Late Trace period. While across-mean behavioral decoding was stronger in the control than the lesioned animals (effect of lesion: *p* = 0.006, 0.001, 0.173 for Nothing vs. Fixed, Nothing vs. Variable, and Fixed vs. Variable, respectively), both groups of animals clearly learned the task and had above-chance across-mean decoding (*p* < 0.001 compared to chance level of 50% for both Nothing vs. Fixed and Nothing vs. Variable in control as well as lesioned animals). Interestingly, Fixed vs. Variable classification was also weakly significant (*p* = 0.032 relative to chance level of 50%) for fully intact control animals, providing behavioral evidence that they did in fact learn this distinction.

**Extended Data Fig. 10.**
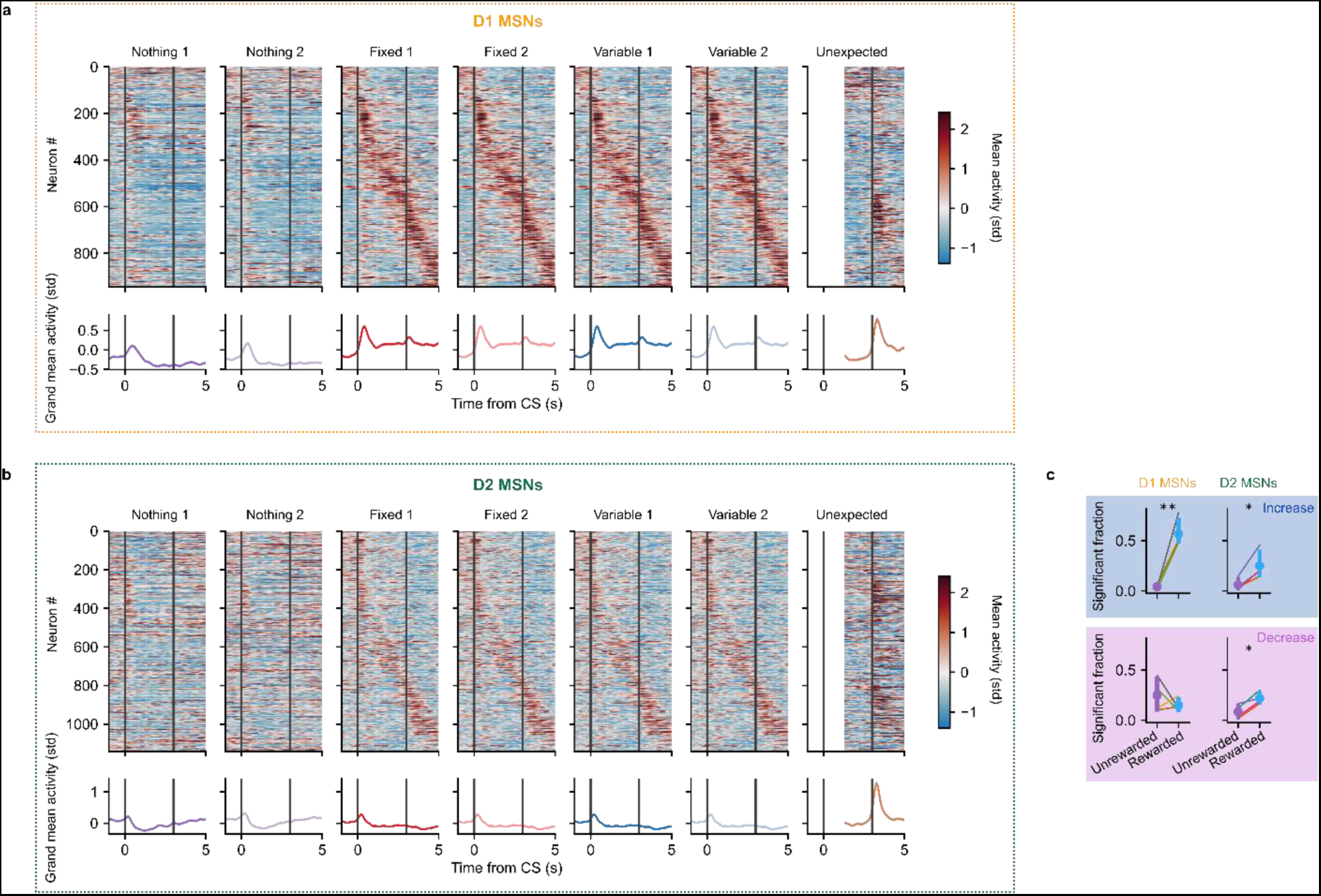
**| Additional data for two-photon calcium imaging. a**, D1 MSN activity. *Top*, heatmaps showing average z-scored deconvolved calcium activity in response to each odor for each neuron, as in Extended Data Fig. 2b. Unexpected reward trials were cropped on the left to include only continuous acquisitions. *Bottom*, grand average z-scored deconvolved calcium activity across all neurons. **b**, Same as **a**, but for D2 MSN activity. **c**, Fraction of neurons whose Late Trace activity increased (*top*) or decreased (*bottom*) relative to Baseline, shown separately for D1 (*left*) and D2 (*right*) MSNs and Unrewarded (Nothing) versus Rewarded (Fixed and Variable) odors (*x*-axis); these trial types were pooled before analysis. As expected, a larger fraction of D1 MSNs increases to Rewarded rather than Unrewarded odors (*p* = 0.006), while there is no difference in the fractions that decrease (*p* = 0.423). Meanwhile, for D2 MSNs, a significantly greater fraction of neurons change their activity on Rewarded compared to Unrewarded trials, by either increasing (*p* = 0.022) or decreasing (*p* = 0.016) their activity relative to Baseline. Asterisks and *p*-values report the results of paired *t*-tests on Rewarded vs. Unrewarded fractions across mice.

**Extended Data Fig. 11.**
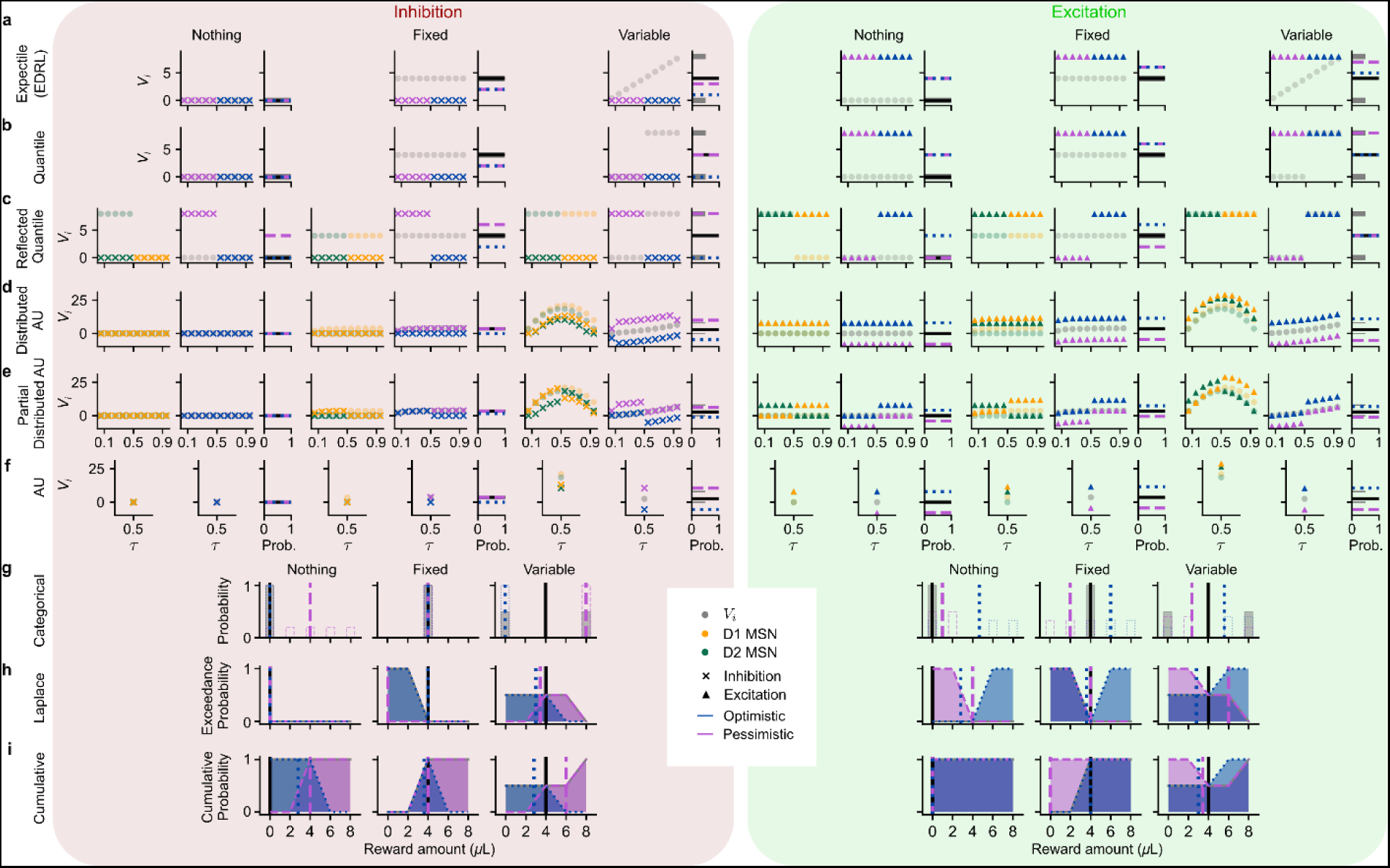
**| Additional detail for distributional model manipulations. a**, Schematic showing how optogenetic perturbations were simulated for an expectile code (from EDRL). Optimistic (blue) or pessimistic (purple) predictors were shifted from their original values (semi-transparent grey circles) and clamped to low or high values to mimic inhibition (*left*, “x”s) or excitation (*right*, triangles), respectively. Panels on the right depict the ground-truth reward distribution, its mean (black line), and the means of the manipulated sets of value predictors (blue or purple dashed lines). **b**, Same as **a**, but for a quantile rather than expectile code. **c**, Same as **b**, but for a reflected quantile code. The additional, leftmost panel for each distribution depicts the activity of D1 (yellow) and D2 (green) MSNs at baseline (semi-transparent circles) and after manipulations (opaque “x”s and triangles). These are what are directly clamped by the simulated optogenetic inhibition or excitation. As a result, the effect on the implied value predictors (middle panel) corresponding to D2 MSNs are of opposite sign, as is the change in predicted mean (right panel). **d**, Same as **c**, but for the Distributed Actor Uncertainty (AU) model. Since D1 and D2 MSN activities in this model can exceed the maximum reward value, the left panel shows that perturbations were simulated by adding or subtracting a fixed amount from each activity level (opaque “x”s and triangles) relative to baseline (semi-transparent circles). The middle panel plots the resulting value predictors, computed as the pointwise differences between D1 and D2 MSN activities, for pessimistic (purple) and optimistic (blue) manipulations in comparison to baseline (grey semi-transparent circles). **e**, Same as **d**, except that only the optimistic or pessimistic half of MSNs were manipulated to simulate perturbations of D1 or D2 MSNs, respectively. **f**, Same as **d**, except for the original Actor Uncertainty (AU) model in which there is only one pair of value predictors with balanced learning rates (*τ* = 0.5). **g**, Schematic showing how optogenetic perturbations were simulated for a categorical code (from CDRL), which effectively represents the reward distribution using a histogram. Pessimistic (0, 2 μL; purple) or optimistic (6, 8 μL; blue) bins were clamped to 0 or 1 to simulate inhibition or excitation, respectively, relative to baseline (grey). The resulting distributions were normalized to sum to one (see Methods). Dashed vertical lines show the means of the ground-truth (black) and manipulated distributions. **h**, Same as **g**, except for a Laplace code^83^ in which each neuron corresponds to the height of 1 - CDF at a particular point. While the baseline case is always monotonically decreasing, simulated excitation or inhibition can change this. Means were computed by differentiating and then normalizing (see Methods). **i**, Same as **h**, except for a cumulative code where each neuron corresponds to the height of the CDF at a particular point.

**Extended Data Fig. 12.**
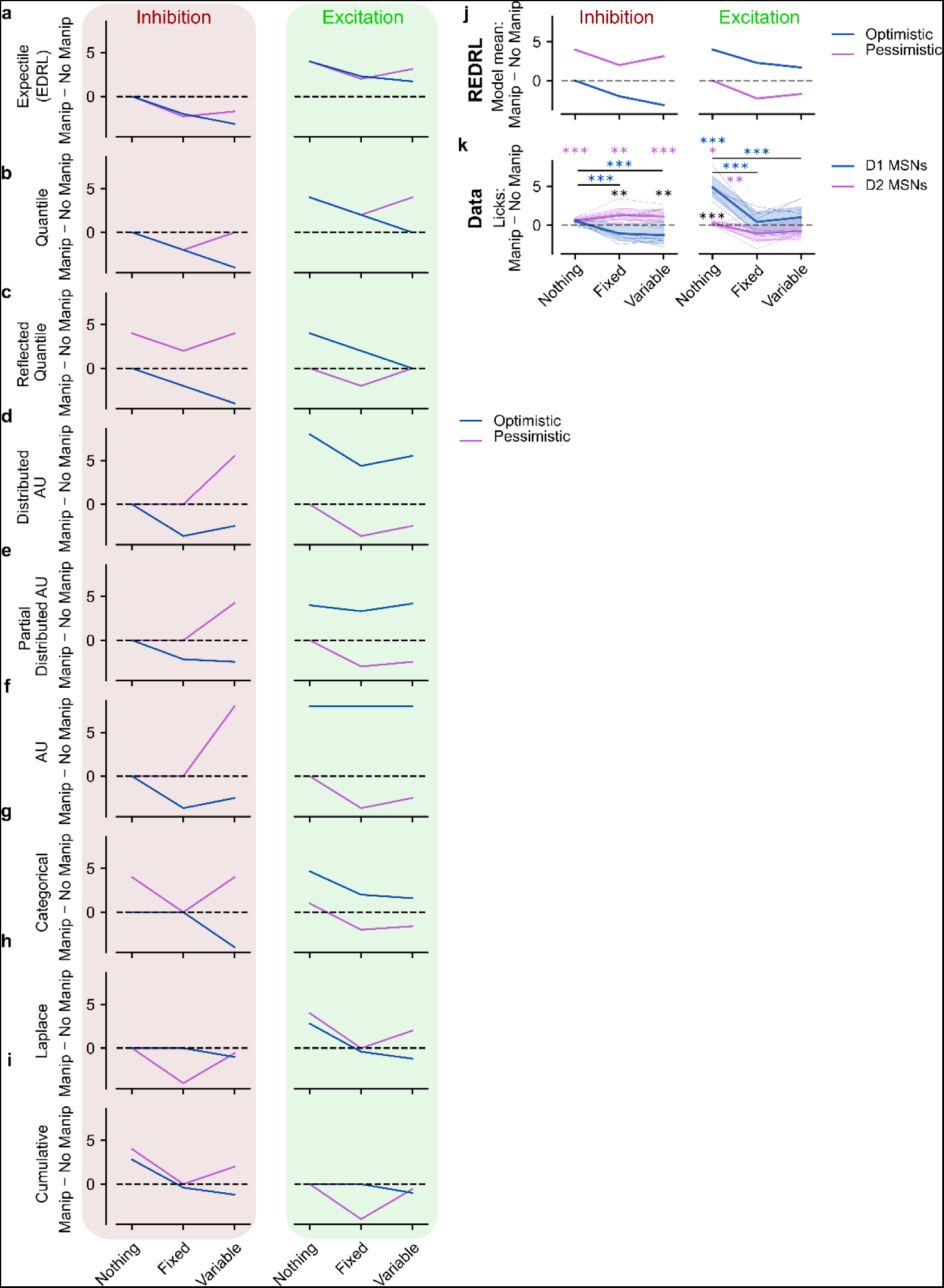
**| Summary of alternative model predictions. a-i**, Predicted difference in mean reward due to inhibition (*left*) and excitation (*right*) for each of the alternative models in Extended Data Fig. 11. **j**, REDRL model predictions for mean reward, copied from Fig. 6e, for comparison. **k**, Actual differences in licking, copied from Fig. 6f, for comparison.

